# Temperature-driven isoform switching reprograms developmental gene expression in a human fungal pathogen

**DOI:** 10.64898/2026.08.26.747414

**Authors:** Murat C. Kalem, Mark Voorhies, Natalie Markman, Anita Sil

## Abstract

Post-transcriptional regulation is key to development, and yet little is known about how RNA isoform choice contributes to developmental choices in fungi. Here we assemble the first isoform-level transcriptome of *Histoplasma*, a ubiquitous human fungal pathogen that grows as an infectious environmental form (hyphae) or a pathogenic host form (yeast) in response to temperature. We find extensive morphology-associated longer leader and trailer isoforms, rapid temperature-driven transcription start site remodeling, and inclusion of regulatory elements in longer leaders. We observe isoform-specific patterns of ribosome and polysome association, indicating that isoform switching regulates the proteome. Our results suggest a model in which isoform diversity and rapid isoform switching are central to post-transcriptional regulation during thermal dimorphism, enabling precise and timely translation of factors needed to establish and maintain hyphal and yeast forms. These studies illuminate how eukaryotic systems utilize post-transcriptional regulation to determine developmental states in response to external stimuli.

## Introduction

Eukaryotic transcriptomes are complex and harbor an impressive level of RNA isoform diversity^1–4^. Multiple RNA isoforms per gene can be generated by alternative transcription start sites (TSSs), splicing, and 3’ end processing^5^. These isoforms are a major source of regulatory diversity in eukaryotes, altering translation, RNA stability and localization^6–13^. The isoform driven post-transcriptional regulation is sometimes achieved through inclusion of upstream/downstream open reading frames (uORFs/dORFs), RNA structural elements and RNA-binding protein (RBP) binding sites in longer 5’ leader and 3’ trailer isoforms^14–21^. RNA isoform diversity plays an important role in development and cellular differentiation^22–25^. For example, short-to-long 5’ leader isoform switching in *Saccharomyces* is implicated in regulating protein levels during meiosis^8,26^. 5’ leader isoform switching controls translation initiation during zebrafish embryogenesis^27^. Likewise, 5’ leader and 3’ trailer isoforms control translation by regulating polysome recruitment during differentiation of human stem cells^28,29^.

As simpler eukaryotes, fungi provide an opportunity to comprehensively study the impact of RNA isoform variation. However, isoform-resolved transcriptome annotations remain scarce in fungi, even for major human pathogens. Recent isoform-resolved annotations in model infectious fungi including the human pathogen *Cryptococcus neoformans* showed that alternative TSS usage and 5’ leader isoforms are common in *C. neoformans,* arise in response to environmental cues and are able to control translation initiation^30,31^. Targeted studies in the plant pathogen *Metarhizium* showed that longer 5’ leader isoforms control precise expression of a protein that controls saprophyte-to-pathogen transition^32^. These studies highlight that isoform biology is consequential but largely remains unmapped in medically important fungi.

The ability of fungi to regulate their development in response to environmental cues makes it possible to expose these organisms to a variety of conditions and assess the consequences on RNA isoform biology. For example, thermally dimorphic fungi such as *Histoplasma ohiense* reprogram their morphology in response to mammalian host temperature to establish disease, making it biologically relevant to assess the production and function of temperature-dependent isoforms^33,34^. *Histoplasma* grows as hyphae in the environment and undergoes a temperature-dependent transition to a yeast form that is thought to be critical to cause disease in mammalian hosts^35,36^. This transition can be recapitulated in the laboratory by shifting culture temperatures^37,38^. To date, the only known regulators of thermal dimorphism include a handful of transcriptional regulators and signaling components, and the post-transcriptional logic of this developmental switch remains enigmatic^34,39–47^.

Prior work defined the gene expression changes between *Histoplasma* yeast and hyphae and identified a small set of genes encoding longer 5’ leader isoforms using short-read RNA-seq^48–50^. This work identified *RYP2*, a transcription factor that is <u>R</u>equired for <u>Y</u>east-<u>P</u>hase growth, as one of the genes that encodes a longer 5’ leader isoform in hyphae at 22°C but not in yeast at 37°C^49^. This initial observation suggested that changes in 5’ leader diversity can contribute to thermal dimorphism by fine-tuning expression of factors involved in this critical developmental switch. However, the global isoform landscape, and whether 5’ leader and 3’ trailer choice is temperature-responsive and post-transcriptionally consequential, remains unknown.

Here we assemble the first isoform-level transcriptome of *Histoplasma ohiense* by integrating long- and short-read RNA-seq approaches. We find extensive morphology-associated longer 5’ leader and 3’ trailer isoforms, rapid TSS remodeling in response to temperature, inclusion of regulatory elements in longer 5’ leaders such as uORFs, and distinct ribosome and polysome association patterns by isoform type. Our findings suggest that alternative 5’ leader and 3’ trailer isoforms provide a regulatory layer for temperature response, thermal dimorphism and pathogenesis. More broadly, resolving isoform regulation in fungi that undergo distinctive developmental transitions illuminates strategies of post-transcriptional control across kingdoms.

## Results

### *Histoplasma* transcriptome assembly reveals a complex isoform-level RNA diversity in yeast and hyphae

To establish *Histoplasma* as an organism to explore temperature-responsive biology and to begin illuminating the molecular principles underlying thermal dimorphism, we built the first isoform-level transcriptome assembly of *Histoplasma* using steady-state yeast and hyphal cells grown at 37°C and 22°C, respectively (**Fig 1A**). We employed an array of complementary sequencing approaches, including Nanopore direct RNA-seq, Illumina paired-end RNA-seq and 5’-seq, which allowed mapping of transcription start sites (TSSs), to capture the full isoform diversity in yeast and hyphae (**Fig 1B**). We clustered TSSs together to establish transcription start region (TSR) annotations requiring each distinct TSR to be at least 25 bases from the next. Illumina reads were used to extract high-resolution splice junction information. Then, we established a computational pipeline to utilize our complementary datasets to build transcript annotations (**Fig 1C**). We established the first version of the isoform annotations using FLAIR (<u>F</u>ull-length <u>a</u>lternative isoforms analysis of <u>R</u>NA) and further filtered isoform annotations based on experimental support (full pipeline is detailed in the Materials & Methods). Briefly, we ensured that the 5’-end annotations from FLAIR agreed with 5’-seq TSR annotations, determined if putative isoform models mapped partially or fully to known gene models, removed low-abundance isoforms, and resolved misannotated or erroneously fused genes. These efforts allowed us to establish a high-confidence isoform-level transcriptome assembly of *Histoplasma*. We then added coding sequence information, PFAM domain annotations, transcription factor annotations, and poly(A) tail information per transcript to further annotate isoform features.

**Figure 1.**
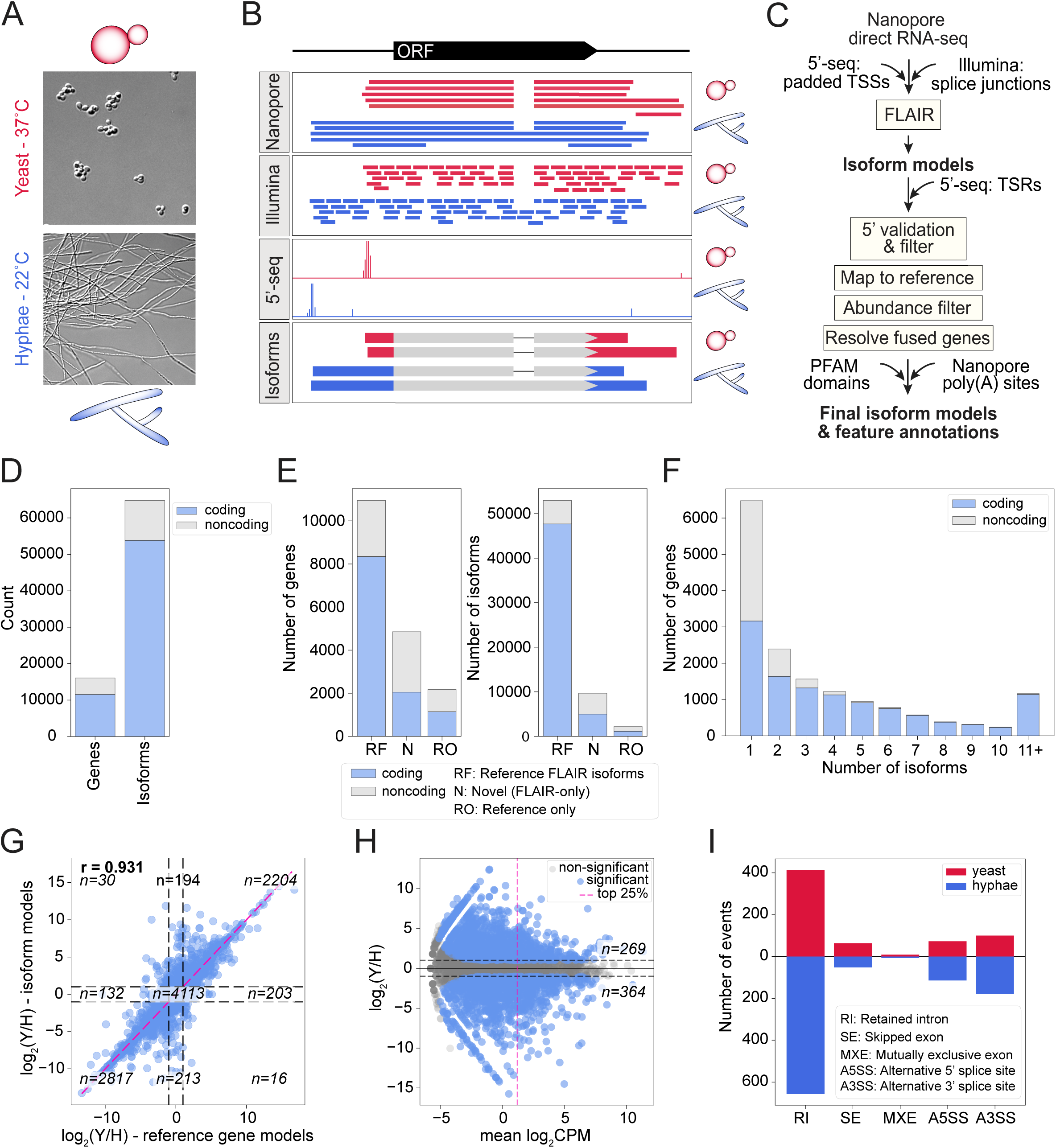
An integrated transcriptome assembly and annotation approach uncovers complex isoform-level RNA diversity in *Histoplasma* yeast and hyphae. A) Representative micrographs show the yeast and hyphal cells at the time of harvest for RNA extraction. B) Integrated RNA-seq approaches utilized for transcriptome assembly and annotation include long-read Nanopore Direct RNA sequencing, short-read paired-end Illumina RNA-seq and 5’-seq to map transcription start sites (TSSs). C) Computational pipeline for transcriptome assembly is shown. Complementary approaches allow for a high-confidence isoform-level assembly. Nanopore reads along with padded TSSs from 5’-seq and splice junctions from Illumina reads were used as inputs to the FLAIR transcriptome assembler. D) Number of coding and non-coding genes and isoforms is shown. E) Number of genes and isoforms in each assembly category is shown. RF: genes/isoforms assembled in our final transcriptome assembly that map to a gene in the reference gene annotations. N: Novel genes/isoforms present in the final transcriptome assembly but are not present in reference. RO: Gene annotations that exist in the reference but are not present in our transcriptome assembly. F) Number of isoforms per coding and noncoding gene is shown. G) Comparison of differential yeast vs. hyphae gene expression quantification from different transcriptome models is shown. Each point is a gene from the RF set. X values are edgeR estimated log_2_(Y/H) contrasts based on kallisto quantification relative to the single-isoform gene models in the reference set. Y values are edgeR estimated log_2_(Y/H) contrasts based on summing, for each gene, all isoform level kallisto estimates relative to our new transcriptome assembly. Dashed horizontal lines are drawn at 2-fold change marks. Pink line represents x=y. Pearson correlation coefficient is highlighted. “n” values indicate number of genes in each of the 9 categories delineated by the 2-fold change marks. H) Scatter plot shows yeast vs. hyphae differential gene expression relative to mean abundance, in units of log_2_ counts per million (log_2_(CPM)). Each point is a Novel gene (N). Horizontal dashed lines indicate 2-fold change. Vertical dashed pink line indicates the top quartile of all genes by mean abundance. Genes are colored blue if they are significantly differential with a 2-fold cutoff and 5% false discovery rate (FDR). Number of genes from the N set that are in the most abundant quartile and significantly yeast or hyphal enriched are indicated. I) Alternative splicing events in yeast and hyphae are shown. Number of events for each splice category was plotted based on relative enrichment in yeast and hyphae as inferred by rMATS. RI: Retained intron; SE: Skipped exon; MXE: Mutually exclusive exon; A5SS: Alternative 5’ splice site; A3SS: Alternative 3’ splice site.

Our final transcriptome atlas includes 16,074 genes and 64,802 isoforms, of which 11,528 genes and 53,805 isoforms were protein-coding and 4,546 genes and 10,997 isoforms were noncoding (**Fig 1D**). Overall, these isoforms were the result of alternative splicing events, alternative transcription start site selection, and alternative transcription termination and 3’ end processing. We further categorized assembled isoforms by mapping to existing gene models. Isoforms that mapped to the reference gene models were categorized as “Reference-FLAIR” (RF); isoforms that are present in our FLAIR-guided assembly but are not present in reference gene models were categorized as “Novel” (N); and gene annotations that exist in reference gene models but not present in our final transcriptome assembly were categorized as “Reference-only” (RO). The majority of the genes and isoforms belonged to the RF category (**Fig 1E**). Unsurprisingly, investigating the abundance of all isoforms in each category revealed that while RF isoforms were generally more abundant, N and RO isoforms had low abundance as indicated by log_2_(CPM) values (CPM: counts per million) **(Fig S1)**. Overall, the majority of the genes (68.15%) had two or more RNA isoforms (**Fig 1F**).

We then evaluated the impact of new isoform models on gene expression quantification in yeast and hyphae. We mapped the Illumina paired-end RNA-seq data to either reference gene models or isoform models that we established in this study and determined yeast vs. hyphae (Y/H) differential expression using edgeR. This revealed that while quantification based on reference and isoform models mostly agree (Pearson’s r = 0.931) (**Fig 1G**), using isoform models improved the quantification, especially where transcript borders were previously not accurately annotated. In addition to fine-tuning gene expression quantification, our new isoform level transcriptome assembly identified a group of novel genes that the reference gene set did not include. Even though most of this novel category was low-abundance transcripts, we identified 269 yeast-enriched and 364 hyphal-enriched novel transcripts where their abundance (log_2_(CPM)) values was in the top 25^th^ quantile and comparable to the abundance of RF transcripts (**Fig 1H, Fig S1).** BLAST analysis of novel transcripts revealed novel genes encoding for ribosomal proteins and small secreted effectors **(Fig S2)**. Many genes in the novel category had already been annotated in other *Histoplasma* species and other fungi, but some did not have a high-confidence BLAST hit. To further establish the isoform level diversity, we cataloged different alternative splicing events using rMATS (replicate multivariate analysis of transcript splicing). This revealed, in total, 659 or 1008 alternative splicing events that were more prevalent in yeast or hyphae, respectively (**Fig 1I**). This comprehensive and well-annotated transcriptome atlas will allow more detailed molecular investigations of gene expression and mechanisms underlying thermal dimorphism in *Histoplasma*.

### Long 5’ leader RNA isoforms exhibit morphology-specific expression and include key regulators of thermal dimorphism

Since 5’ leaders and 3’ trailers harbor important post-transcriptional regulatory elements and meaningfully influence translation and RNA decay, we set out to determine the 5’ leader diversity in the *Histoplasma* transcriptome. We first evaluated the 5’ leader length in all protein coding RNA isoforms. The median and mean 5’ leader length were 232 nt and 365 nt, respectively (**Fig 2A**). *Histoplasma* 5’ leader lengths were more comparable to those of *Homo sapiens* and *Drosophila melanogaster*^51^ (median length: 218 nt and 186, nt respectively) than other fungi (median length: *Saccharomyces cerevisiae*^51–53:^ 53 nt, *Cryptococcus neoformans*^54^: 105 nt, *Aspergillus fumigatus*^55^: 49 nt, *Coprinopsis cinerea*^56^: 68 nt), suggesting a more complex regulation.

**Figure 2.**
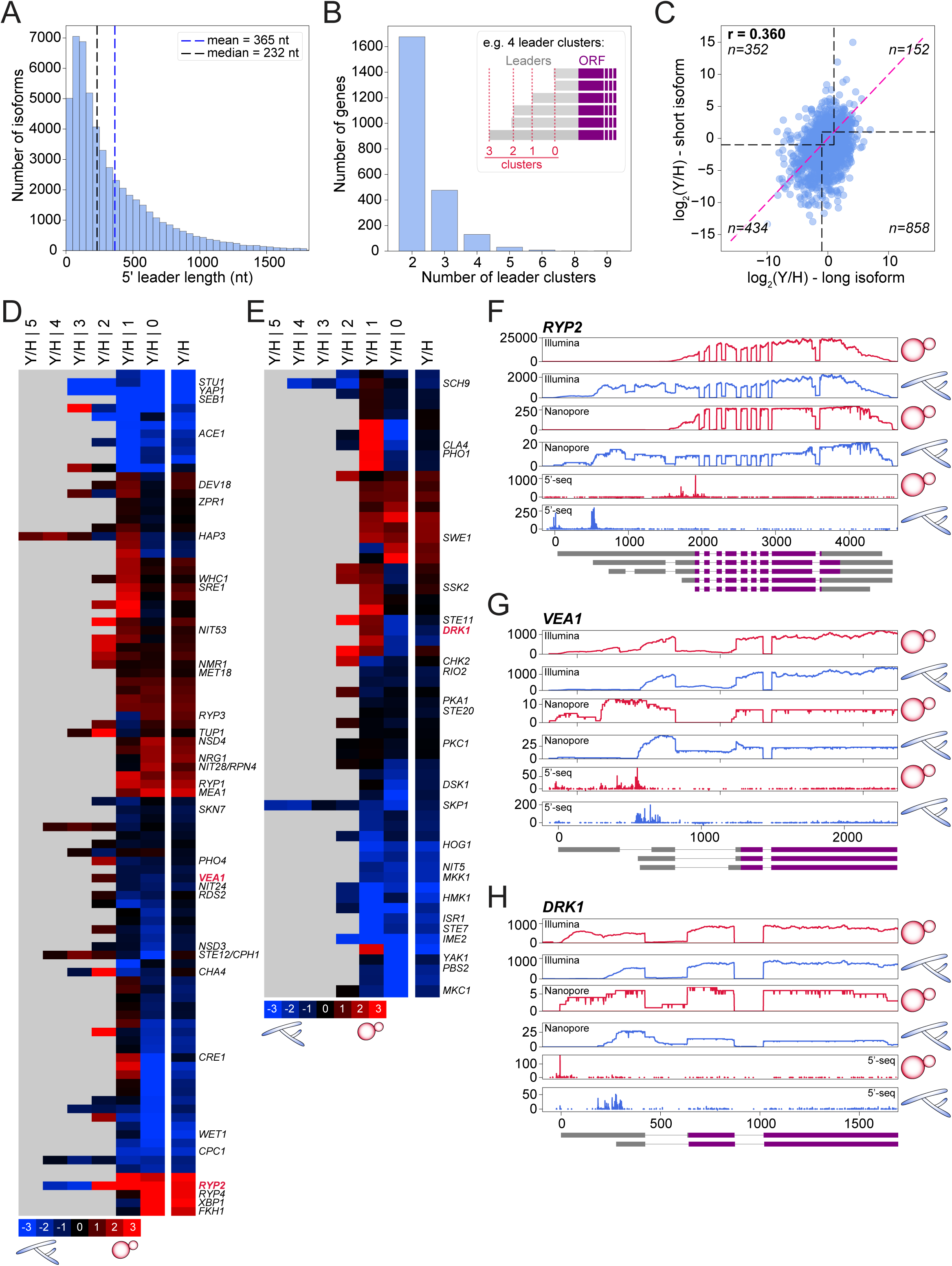
Long 5’ leader RNA isoforms with morphology-specific expression include key regulators of thermal dimorphism. A) Distribution of 5’ leader lengths is shown. Isoforms with leader lengths above the 99^th^ percentile (538 isoforms) are excluded from the plot. Mean and median leader lengths are indicated by blue and black dashed lines, respectively. B) Number of genes per longer leader cluster is shown. For each gene, 5’ leader sequences across all isoforms are clustered by start position, with clusters separated by a minimum of 25 bases. The inset shows a representative clustering scheme. Clusters are ranked in an ascending order, with rank 0 assigned to the shortest leader cluster. C) Scatter plot shows yeast vs. hyphae differential expression of short and long 5’ leader isoforms. For simplicity, the rank 0 cluster was designated as “short”, and the remaining clusters were designated as “long”. X and Y values were edgeR-estimated log_2_(Y/H) contrasts for long and short isoforms, respectively. Dashed horizontal lines are drawn at 2-fold change marks. Pink line represents x=y. Pearson correlation coefficient is highlighted. “n” values indicate the number of genes in each of the 4 categories delineated by the 2-fold change marks. Top-right and bottom-left quadrants indicate genes where both long and short leader isoform expression change in the same direction. Bottom right and top left quadrants indicate genes where longer leader expression is more abundant in yeast or hyphae, respectively, while the short isoform expression is either unchanged or their expression change in the opposite direction. D) Transcription factor-encoding genes with longer 5’ leader isoforms are shown. Heatmap showing the log_2_ fold-change (yeast relative to hyphae) for the overall gene expression (Y/H) and for each 5’ leader cluster ranked by start position (Y/H | 0 to 5), where 0 represents the shortest and 5 the longest cluster. Gray indicates no data. E) Heatmap, as in D, shows protein kinase encoding genes. F) *RYP2,* G) *VEA1, and* H) *DRK1* read coverage tracks are plotted. Plots for *VEA1* and *DRK1* focus on the 5’ region. From top to bottom: Illumina, Nanopore, and 5’-seq read coverage in yeast and hyphae. The x-axis indicates relative genome location in bases and the y-axis indicates number of reads. Representative isoform models are included at the bottom with introns as lines and exons as rectangles. 5’ leaders and 3’ trailers are shown as gray rectangles; and coding sequences (CDS) are shown as purple rectangles.

We then explored the transcripts that express both a long and a short 5’ leader isoform, since presence of both for a given gene implies that the organism has the possibility to switch between different isoforms depending on its biological state. Long and short isoform switching has been documented in fungi including *Saccharomyces* and *Histoplasma*. In *Saccharomyces*, long 5’ leader isoforms that are poorly translated, also referred to as long undecoded transcript isoforms or LUTIs, were characterized during meiosis and are often associated with poor translation and repression of transcription from proximal promoters^8,26,57,58^. In *Histoplasma*, we previously identified 187 genes with longer 5’ leader isoforms by using paired-end Illumina RNA-seq and only considering longer leader isoforms that were observed in 3 out of 4 *Histoplasma* strains^49^. Here we analyzed the prevalence of longer 5’ leader isoforms by clustering 5’ start sites of all isoforms of a given gene that were within 25 bases from each other and designating each set with a consecutive number (**Fig 2B, inset)**. We indexed the shortest cluster as 0 and cluster 1, for example, indicated the next longest cluster that was at least 25 bases from cluster 0. We identified 2,328 genes with a longer 5’ leader isoform (**Fig 2B**). Most of these genes had only two 5’ leader clusters. We then determined the expression of long and short 5’ leader isoform clusters in yeast vs hyphae. For simplicity, the cluster 0 was designated as ‘short’ and the rest of the clusters were collapsed and designated as ‘long’. This comparison was restricted to genes with significant differential expression of the long and/or short isoform groups (FDR *≤* 0.05, *n* = 1,796). Among these genes, Y/H differential expression of long or short isoforms was only weakly correlated (Pearson’s r = 0.360) (**Fig 2C),** meaning that long and short isoforms of a given gene often did not show the same Y/H differential expression. We identified 152 and 434 genes where both long and short isoform groups were more enriched in yeast or hyphae, respectively. The discordant quadrants contained genes in which the long and short isoforms are differentially expressed in opposite directions with respect to Y and H, or in which one isoform group (short or long) is relatively unchanged (|log_2_(Y/H)|<1) while the other is differentially expressed.

To determine if genes that encode longer leader isoforms were enriched for any particular biological function, we used InterPro domain annotations and their corresponding gene ontology associations to broadly evaluate gene ontology terms. This revealed two important themes: transcription regulation (GO:0006355, n_query/n_total: 75/173; GO:0000981, 46/111; GO:0003700, 24/51) and protein kinase activity (GO:0004672, n_query/n_total: 60/157). We updated the list of transcription factors and kinases and included those that were not captured in the InterPro and GO approach. Yeast versus hyphae differential expression of isoforms in each leader cluster (Y/H | cluster number) along with overall gene level expression (Y/H) are shown for all genes that fall under transcription regulation (**Fig 2D**) and protein kinase activity GO terms (**Fig 2E**).

Transcription factors with longer 5’ leaders included key regulators of thermal dimorphism in *Histoplasma*, such as Stu1^44^, Wet1^42^, Ryp1-4 (Ryp1, Ryp2, Ryp3, Ryp4)^39–41^, and Vea1^47^ (**Fig 2D**). All of these transcription factors have been previously shown to control thermal dimorphism, either promoting yeast or hyphal morphology. We noted that the expression of the longer 5’ leader isoforms was not always differential between yeast and hyphae. Some differential longer leader cases showed increased expression in yeast and some in hyphae. For example, while *RYP2* had a longer 5’ leader isoform in hyphae, which is supported by Illumina, Nanopore and 5’-seq datasets, *VEA1* had a longer 5’ leader isoform in yeast (**Fig 2F, 2G)**.

Kinases with longer 5’ leader isoforms likewise included regulators of thermal dimorphism and stress signaling. One intriguing example is the histidine kinase Drk1 (ortholog of *C. albicans* Nik1), which is an important regulator of thermal dimorphism in *Blastomyces* and *Histoplasma*^45^. In *Histoplasma*, Drk1 is necessary for yeast morphology and virulence^45^. We found that *DRK1* mRNA had a longer leader isoform in yeast (**Fig 2H**). Intriguingly, mRNAs encoding orthologs of proteins that function in the Drk1 pathway in other fungi also displayed long and short 5’ leader isoforms in *Histoplasma*. RNAs encoding phosphotransfer protein Ypd1 and the response regulator Ssk1 had longer 5’ leaders in yeast, whereas MAP kinases Ssk2 (MAPKKK) and Hog1 (MAPK) had longer 5’ leaders in hyphae. In contrast, Pbs2 (MAPKK) mRNA had a short 5’ leader in yeast. This enrichment of longer leaders for a known signaling pathway was surprising given the low prevalence of longer leaders overall, and suggests potential post-transcriptional regulation of this pathway.

### mRNAs encoding transcription factors and RNA-binding proteins have long 3’ trailer RNA isoforms with morphology-specific expression

Like 5’ leaders, 3’ trailers can regulate post-transcriptional processes including translation and RNA decay. Our transcriptome annotation approach, which included Nanopore direct RNA-seq of polyadenylated mRNAs, allowed us to explore 3’ trailer diversity in *Histoplasma*. We first evaluated the 3’ trailer length in all protein coding RNA isoforms. The median and mean 3’ trailer length were 364 nt and 470 nt respectively (**Fig 3A**). The median 3’ trailer length in *Histoplasma* exceeded reported values for other fungi such as *S. cerevisiae*^59^ (121 nt), *C. neoformans*^54^ (127 nt), *A. fumigatus*^55^ (184 nt), and the fruiting multicellular mushroom *C. cinerea*^56^ (141 nt) as well as the higher eukaryotes *D. melanogaster*^60^ (224 nt) and *Caenorhabditis elegans*^61^ (130 nt), but was shorter than that of *H. sapiens*^62,63^ (1,200 nt). Thus, *Histoplasma* RNAs have unusually long 3’ trailers relative to other fungi examined.

**Figure 3.**
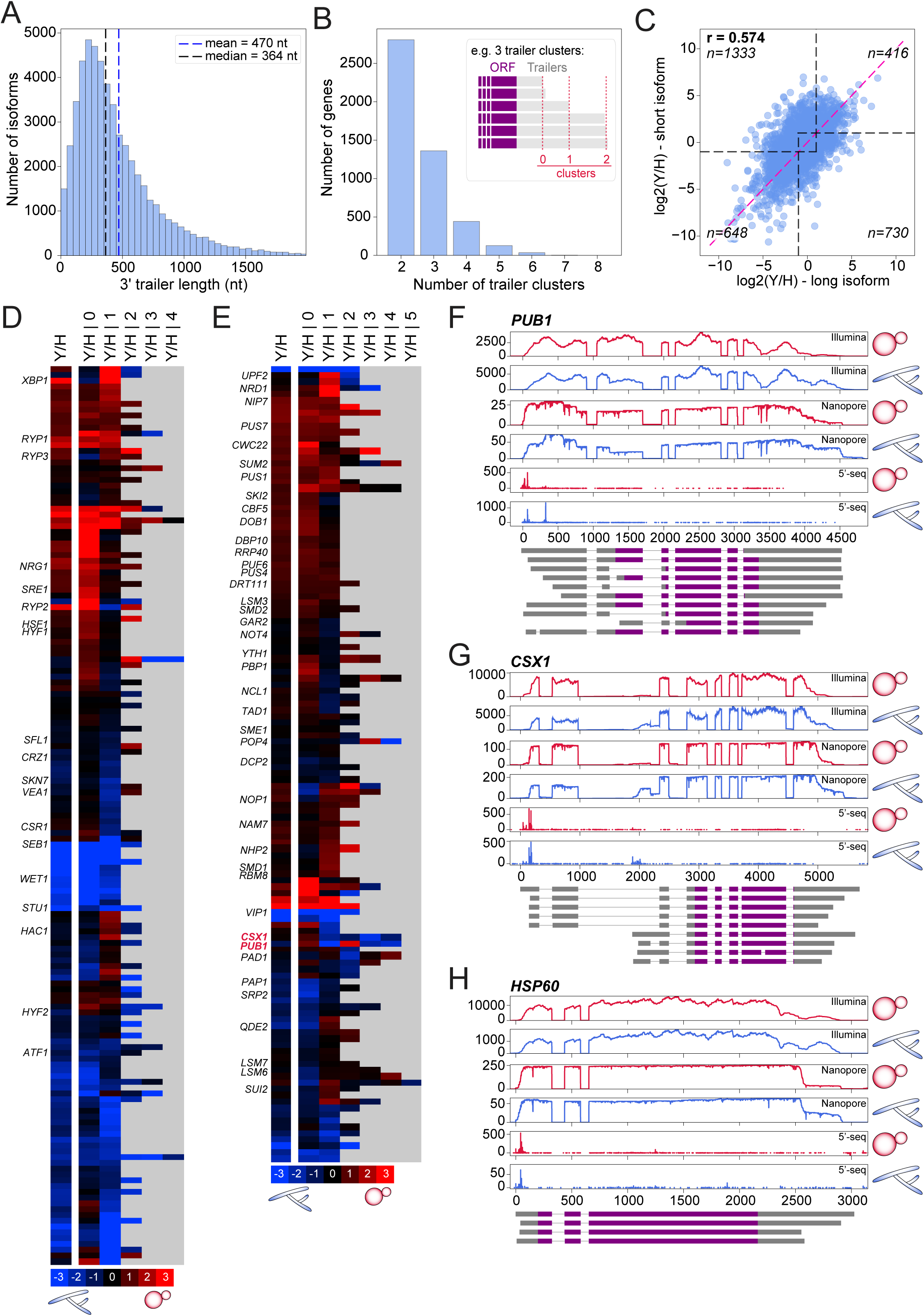
Long 3’ trailer RNA isoforms with morphology-specific expression include transcription factors and RNA-binding proteins. A) Distribution of 3’ trailer lengths is shown. Isoforms with trailer lengths above the 99^th^ percentile (538 isoforms) are excluded from the plot. Mean and median trailer lengths are indicated by blue and black dashed lines, respectively. B) Number of genes per longer trailer cluster is shown. For each gene, 3’ trailer sequences across all isoforms are clustered by start position, with clusters separated by a minimum of 25 bases. Clusters are ranked in an ascending order, with rank 0 assigned to the shortest trailer cluster. C) Scatter plot shows yeast vs. hyphae differential expression of short and long 3’ trailer isoforms. For simplicity, the rank 0 cluster is designated as short, and the remaining clusters are designated as long. X and Y values are edgeR-estimated log_2_(Y/H) contrasts for long and short isoforms, respectively. Dashed horizontal lines are drawn at 2-fold change marks. Pink line represents x=y. Pearson correlation coefficient is highlighted. “n” values indicate the number of genes in each of the 4 categories delineated by the 2-fold change marks. Top-right and bottom-left quadrants indicate genes where both long and short trailer isoform expression change in the same direction. Bottom right and top left quadrants indicate genes where longer trailer expression is more abundant in yeast or hyphae, respectively, while the short isoform expression is either unchanged or changes in the opposite direction. D) Transcription factor-encoding genes with longer 3’ trailer isoforms are shown. Heatmap showing the log_2_ fold-change (yeast relative to hyphae) for the overall gene expression (Y/H) and for each 3’ trailer cluster ranked by start position (Y/H | 0 to 5), where 0 represents the shortest and 5 the longest cluster. Gray indicates no data. E) Heatmap, as in D, shows RNA binding protein encoding genes. F) *PUB1,* G) *CSX1, and* H) *HSP60* read coverage tracks are plotted. From top to bottom: Illumina, Nanopore, and 5’-seq read coverage in yeast and hyphae. The x-axis indicates relative distances in bases and the y-axis indicates number of reads. Representative isoform models are included at the bottom. 5’ leaders and 3’ trailers are shown as gray rectangles; and coding sequences (CDS) are shown as purple rectangles.

We analyzed the prevalence of longer 3’ trailer isoforms by clustering 3’ end positions of all isoforms of a given gene that are within 25 bases from each other and designated each set with a number (**Fig 3B, inset)**. We indexed the shortest cluster as 0 and increasing numbers indicate the next longest cluster that was 25 bases from cluster 0. Like 5’ leaders, presence of both long and short 3’ trailers for a given gene implies that the organism has the possibility to switch between different isoforms depending on its biological state. It has been shown that long and short 3’ leader switching controls the interactions with effectors such as RNA-binding proteins and micro RNAs. We identified 4,780 genes with long and short 3’ trailer isoforms (**Fig 3B**). Most of these genes had only two 3’ trailer clusters. We then determined the expression of long and short 3’ trailer isoform clusters in yeast compared to hyphae (Y/H). For simplicity, the cluster 0 was designated as ‘short’ and the rest of the clusters were collapsed and designated as ‘long’. This comparison was restricted to genes with significant differential expression of the long and/or short isoform groups in Y/H differential expression (FDR ≤ 0.05, n = 4,542). Among these genes, Y/H differential expression of long and short isoforms was moderately correlated (Pearson’s r = 0.570) (**Fig 3C**). We identified 416 and 648 genes where both long and short isoform groups were more enriched in yeast and hyphae, respectively. The discordant quadrants contained genes in which the long and short isoforms change in opposite directions, or in which one isoform group (short or long) was relatively unchanged (|log_2_(Y/H)|<1) while the other was differentially expressed.

We observed that most of the genes encoding transcription factors or transcription regulators with longer 5’ leaders also had longer 3’ trailer isoforms. In fact, out of 2,328 genes with longer 5’ leader isoforms, 1,784 also had longer 3’ trailer isoforms (**Fig S4A**). There was a moderate correlation between the Y/H differential expression of longer 5’ leader and longer 3’ trailer isoforms **(Fig S4B, Pearson’s r = 0.69)**. We noted that the gene ontology terms in the longer 3’ trailer isoform gene set was enriched for RNA binding proteins (RBPs) (GO:0003723, n_query/n_total: 143/189). We then examined yeast versus hyphal differential expression of isoforms in each 3’ trailer cluster (Y/H | cluster number) along with overall gene level expression (Y/H) or all genes that are either involved in transcription regulation (**Fig 3D**) or RNA-binding proteins. (**Fig 3E**). Transcription factors with longer trailer isoforms included critical regulators of thermal dimorphism such as Ryp1, Ryp2, Ryp3, Wet1 and Vea1 (**Fig 3D and Fig 1F**). Additionally, we noted genes encoding heat shock factor Hsf1 and calcineurin-responsive zinc-finger protein Crz1 in our analysis of longer trailer isoforms. Longer 3’ trailer isoforms were found in genes encoding RBPs that mediate critical post-transcriptional processes, including the poly(A) binding protein Pub1 and the RBP Csx1 (**Fig 3F, 3G)**. In *Schizosaccharomyces pombe,* Csx1 regulates gene expression during oxidative stress and sexual differentiation^64^. Both *PUB1* and *CSX1* had short 5’ leader isoforms and long 3’ trailer isoforms that were more abundant in the hyphal morphology. Finally, we identified several heat-shock protein-encoding genes with longer 3’ trailer isoforms including *HSP10*, *HSP60*, *HSP70*, *BiP*, *SSZ1*, and *HSP82*. Longer 3’ trailer isoforms of *HSP60* were more prevalent in hyphae than yeast (**Fig 3H**). Together, these analyses establish extensive 3’ trailer diversity in *Histoplasma* and show that longer 3’ trailer isoforms are found in genes encoding key transcription factors, post-transcriptional regulators and heat-shock proteins.

### Long and short 5’ leader switching occurs rapidly in response to temperature

Since some genes show differential 5’ leader isoforms in steady-state yeast (at 37°C) versus steady-state hyphae (at 22°C), it was interesting to determine if the choice of leader sequence was determined by growth temperature, which would trigger a rapid switch in leader sequence after temperature shift, or cell morphology, which takes days to manifest after temperature shift. We examined the temporal dynamics of 5’ leader isoform switching post-temperature shift from 37°C to 22°C. The TSSs were profiled using 5’-seq at 0, 2, 6 hours, and day 1 and 4 post-temperature shift (**Fig 4A**). We observed that by day 4, cultures were a mixed population of yeast and pseudohyphal cells (**Fig 4A**) that would standardly form hyphae by day 7 at 22°C, similar to our steady-state hyphal samples. To quantify 5’ leader isoform dynamics throughout this time course, TSSs from 5’-seq data were clustered into TSRs and the differential expression profile was determined for each TSR relative to the 0 hour time point. Because individual TSRs from the same gene can have distinct temporal expression profiles, we used principal component analysis (PCA) of TSR expression profiles per gene to classify genes by TSR expression complexity (see Materials & Methods). We restricted this analysis to genes whose expression level, inferred by expression of all TSRs of that gene, was differential at one or more time points. (**Fig 4B, right)**. We identified 891 genes where at least two TSRs changed expression in opposite directions relative to time 0. We referred to this category of genes as class 1. TSRs were differentially expressed either in the same direction as the entire gene (orange in **Fig 4B, left**), or in the opposite direction (purple in **Fig 4B, left**). Some genes showed distal TSRs that tracked with the gene-level TSR expression profile, whereas for others only the more proximal TSRs did, indicating that TSRs giving rise to short or long 5’ leader isoforms can be differentially regulated after an acute temperature shift from 37°C to 22°C. We also observed 672 genes with more than two distinct TSR expression profiles where multiple principal components were needed to capture the temporal TSR expression variance (class 2) **(Fig S5)**.

**Figure 4.**
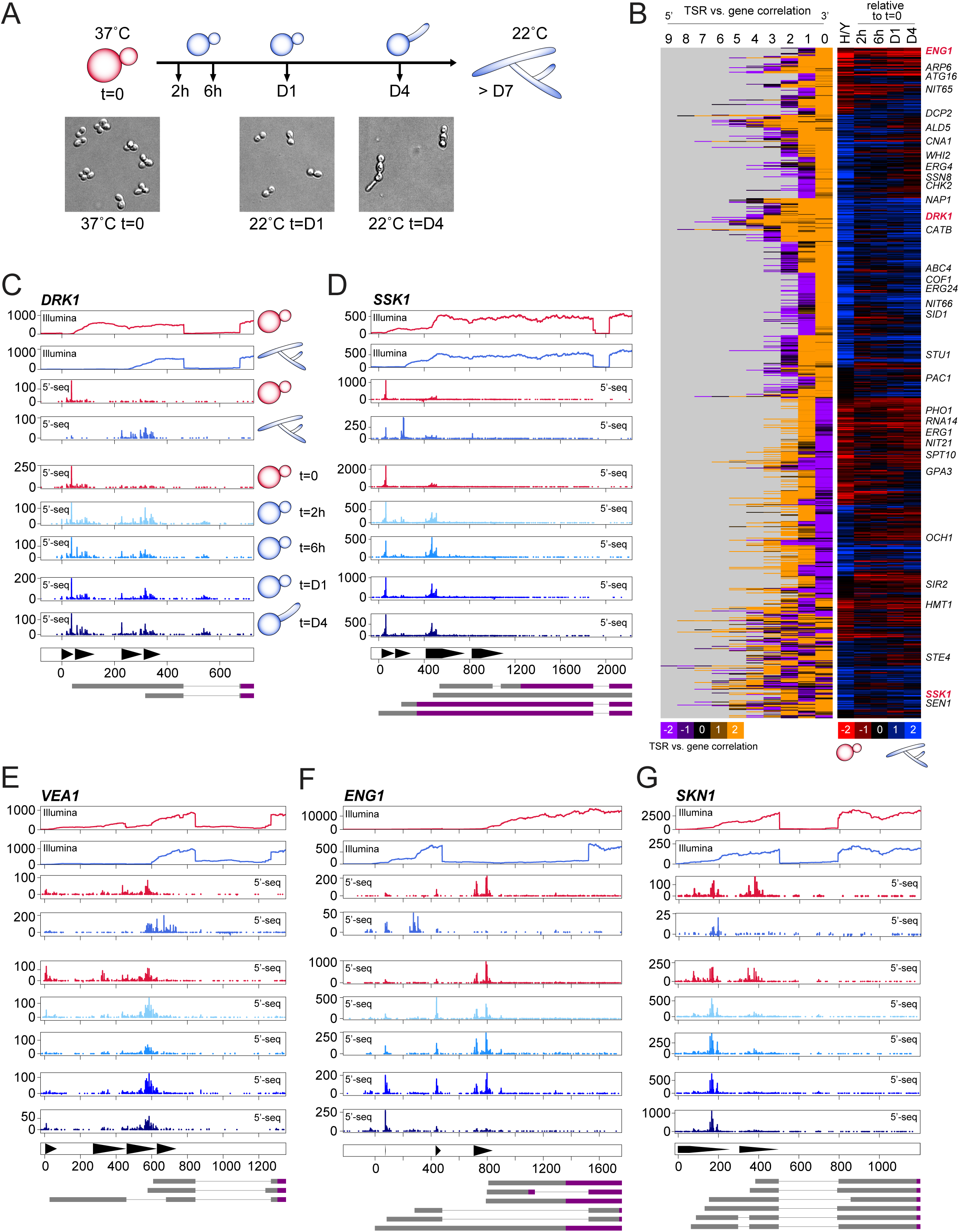
Long and short 5’ leader isoform switching is rapidly responsive to temperature. A) Outline of acute temperature shift time course is shown. Mid-log yeast cells at 37°C were transitioned to 22°C and cells were collected at 2 hours, 6 hours, 24 hours (day 1 - D1), and 96 hours (day 4 - D4) post-temperature shift for RNA extraction and 5’-seq to map temperature-responsive transcriptions start sites and regions (TSSs and TSRs). B) Heatmaps show genes with at least two temperature responsive TSRs where only one principal component (PC1) explains more than 20% of the variance across TSRs of a given gene (class 1 genes). *Right panel:* Differential expression of all summed TSRs of a given gene in steady state hyphae relative to yeast (H/Y), or relative to mid-log yeast at 37°C (t=0) across time points. *Left panel:* PC1 weights on each TSR. Genes shown have at least two TSRs with opposite-sign weights, indicating divergent temperature responsive expression. PCs are oriented so that positive weight (orange) indicates correlation with the gene level expression pattern and negative weight (purple) indicates anticorrelation. TSR 0 denotes the most 3’ TSR and TSR 9 denotes the most 5’ TSR relative to the start codon. Rows are clustered by 5’ to 3’ ordering of PC1 weights on TSRs, then by gene-level expression pattern. Gray indicates no data. C) *DRK1* 5’ region read coverage is plotted. From top to bottom: Illumina and 5’-seq read coverage in yeast and hyphae followed by time course tracks at t=0, 2h, 6h, D1, and D4 post-temperature shift. The x-axis indicates relative distances in bases. Representative isoform models are included at the bottom. 5’ leaders and 3’ trailers are shown as gray rectangles; and CDSs are shown as purple rectangles. D) *SSK1*, E) *VEA1*, F) *ENG1*, and G) *SKN1* read coverage tracks showing the 5’ region are plotted, as in C.

Our systemic classification of temporal TSR expression profiles revealed several genes encoding regulators of thermal dimorphism. *DRK1* had a dominant long 5’ leader in steady-state yeast at 37°C and a dominant short 5’ leader in steady-state hyphae at 22°C (**Fig 4C, top four tracks)**. After the shift from 37°C to 22°C, *DRK1* long leader isoform expression decreased by 2 hours with a concomitant increase in the short leader isoform (**Fig 4C**). *SSK1* likewise had a longer 5’ leader isoform in steady-state yeast and a shorter 5’ leader isoform in hyphae (**Fig 4D, top four tracks)**. A minor TSR producing the shortest *SSK1* 5’ leader was present in steady-state yeast and hyphae but became more abundant by 2 hours post-temperature shift (**Fig 4D**). We noted that utilization of this TSR, which was more abundant during the transition state, would result in a truncated protein product. For *VEA1*, expression from the TSR corresponding to the longest 5’ leader decreased by 2 hours post-temperature shift (**Fig 4E**). We also observed dynamic 5’ leader switching at cell-wall metabolism genes: *ENG1*, which encodes a β-(1,3)-glucanase, had shorter 5’ leader isoforms in yeast and longer 5’ leader isoforms in hyphae (**Fig 4F, top four tracks)**. Longer leader expression increased by 2 hours after temperature shift; we also observed an intermediate-length long leader isoform most prominent in the transition state but with insufficient signal to resolve as an independent component (**Fig 4F**). *SKN1*, which encodes a β-1,6-glucan synthesis protein, expressed both long and short 5’ leader isoforms in yeast but predominantly the long 5’ leader in hyphae (**Fig 4G, top four tracks)**. Short 5’ leader expression of *SKN1* decreased sharply after 2 hours post-temperature shift, while the long 5’ leader was maintained, implying independent regulation of the two TSRs. Together, these examples indicate that long and short 5’ leader isoform usage of key morphology and cell wall genes can change within 2 hours of temperature shift, including transition-specific TSR usage that is not observed in steady-state yeast or hyphae. These rapid changes in TSS selection, and 5’ leader diversity, raise the possibility that early isoform switching contributes to the temperature response during establishment of thermal dimorphism.

### Switching between long and short 5’ leader isoforms determines inclusion of regulatory RNA elements including inhibitory uORFs

5’ leaders can harbor a diverse set of regulatory elements that control gene expression post-transcriptionally, including RBP binding sites, RNA modification sites, internal ribosomal entry sites, RNA structures, upstream AUGs (uAUGs) and uORFs^18–20,51^. Therefore switching between different 5’ leader isoforms can determine if these regulatory elements are included. We asked whether long and short 5’ leader switching introduces uORFs that are translated and associated with reduced CDS translation. uORFs are known to be important in regulating translation under stress conditions^65,66^. We first computationally annotated all possible uORFs and used our published ribosome profiling data to quantify uORF and CDS translation efficiency (TE) (**Fig 5A**). Our new isoform-level transcriptome annotations gave us a higher-resolution framework for interpreting our previously published ribosome profiling data. Translational efficiency (TE) was defined as ribosome footprint signal divided by RNA-seq coverage across uORF or CDS coordinates. Among 19,333 uORFs with sufficient read coverage, 5,044 and 1,402 had more than 2-fold higher TE relative in either yeast or hyphae, respectively. Concurrently, we enriched for cases consistent with uORF-dependent repression of downstream CDS translation by requiring a lower CDS TE. We then identified genes that encode at least one isoform with and one without the uORF, and required expression of the uORF-containing isoform to be consistent with the morphology in which it was repressive. This yielded 24 yeast and 31 hyphal genes with long and short 5’ leader isoforms in which long 5’ leader uORFs were preferentially translated while CDS TE was reduced (**Fig 5B, 5C)**. For example, the long *RYP2* 5’ leader, which was more abundant in hyphae, introduced uORFs with ribosome footprint signal in the 5’ leader and had reduced CDS translation in hyphae (**Fig 5D**). Conversely, a cytochrome P450 gene (ACKS0A_10348) had longer 5’ leaders in yeast whose uORFs were associated with reduced CDS translation in yeast (**Fig 5E**). Together, our data indicate that switching between long and short 5’ leader isoforms can determine inclusion of regulatory elements such as uORFs and is associated with morphology-specific differences in translation.

**Figure 5.**
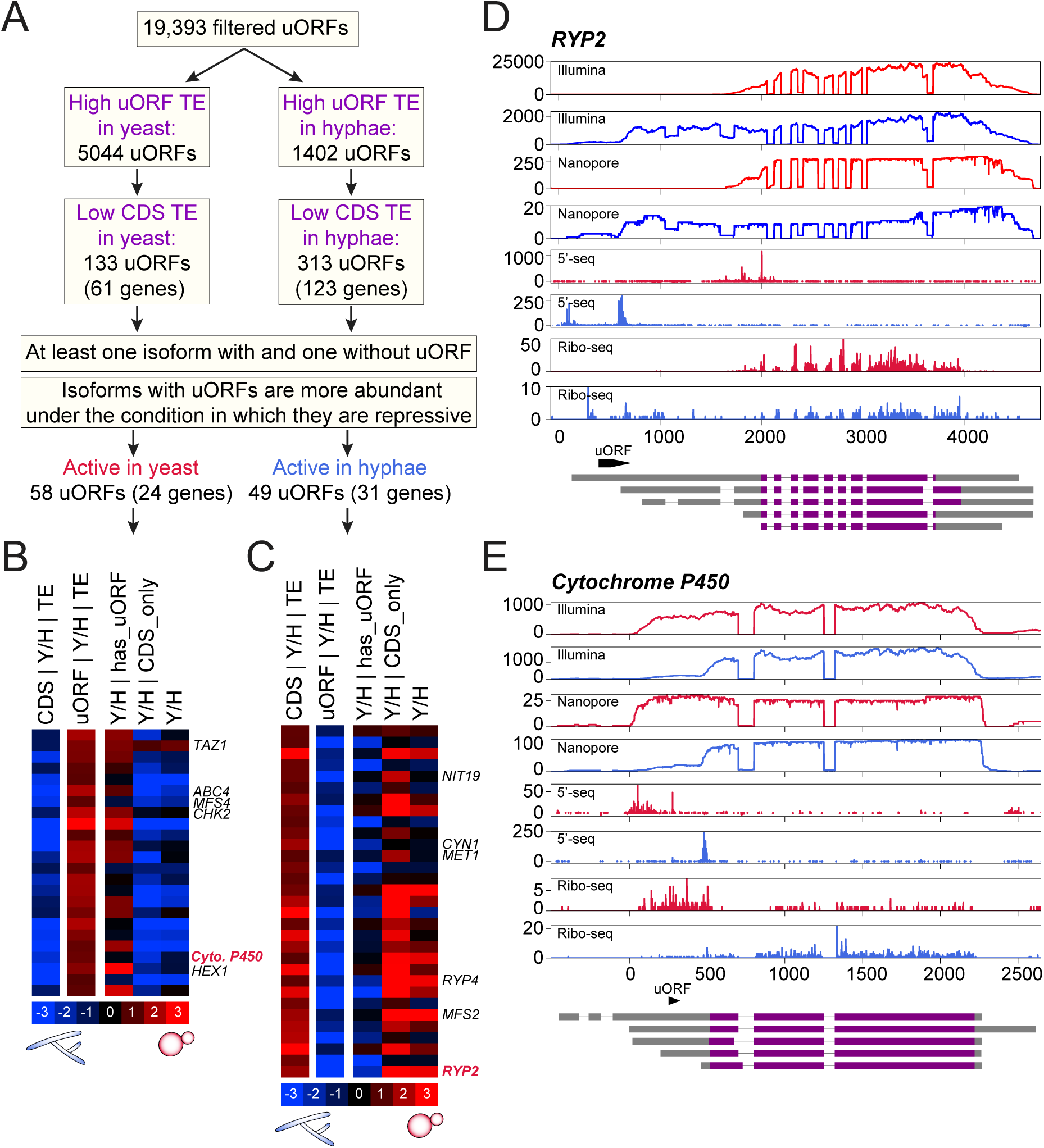
Switching between long and short 5’ leader isoforms determines inclusion of inhibitory uORFs Regulation of inhibitory uORFs is achieved by switching between long and short 5’ leader isoforms. A) Analysis pipeline to determine uORFs in long 5’ leader isoforms is shown. All possible uORFs were annotated and translational efficiency (TE) across this uORF was calculated using Ribo-seq data from Gilmore et al., 2015. Isoforms with high uORF TE and low CDS TE were determined. Of these, genes encoding at least one isoform with and one isoform without uORF was selected along with an expression filter. B) Heatmap shows genes with longer leader isoform-specific uORFs that have ribosome footprints (therefore referred to as active) and correlate with translational repression of the main ORF in yeast. Columns: *Left to right*. Differential TE between yeast and hyphae in the CDS or uORF, differential transcript levels in yeast relative to hyphae for isoforms containing the uORF, isoforms containing the CDS but lacking the uORF, or all isoforms of the gene. C) Heatmap shows genes with longer leader isoform-specific uORFs that are active and correlate with translational repression of the main ORF in hyphae. Heatmap columns are as in B. D) *RYP2,* and D) *Cytochrome P450* read coverage is plotted, as in 2F but adding Ribo-seq data. Ribo-seq data is plotted as read counts per inferred ribosome P-site location.

### Long 5’ leader and 3’ trailer isoforms correlate with distinct ribosome and polysome association

Long and short 5’ leader and 3’ trailer isoforms can affect the inclusion of regulatory RNA elements and thereby influence translation. To investigate isoform-level ribosome association, we performed polysome profiling coupled with RNA-seq in steady-state yeast and hyphal cells. Lysates were separated on a sucrose gradient and fractionated, and RNA from 80S monosome, low-polysome (2-4 ribosomes), and high-polysome (≥5 ribosomes) fractions, along with input, was analyzed by Illumina RNA-seq (**Fig 6A**). We quantified isoform enrichment in polysomes (2 to ≥5 ribosomes) relative to monosomes (*P*) and in ribosome associated RNA (monosome + polysome) relative to input (*R*). Differential *P* and *R* values between yeast and hyphae (*dP* and *dR*) were then used to assess morphology-dependent differences in isoform-level ribosome and polysome association (**Fig 6A**) (see Materials & Methods). We used ribosome and polysome association metrics as a proxy for evaluating isoform-level translation efficiency.

**Figure 6.**
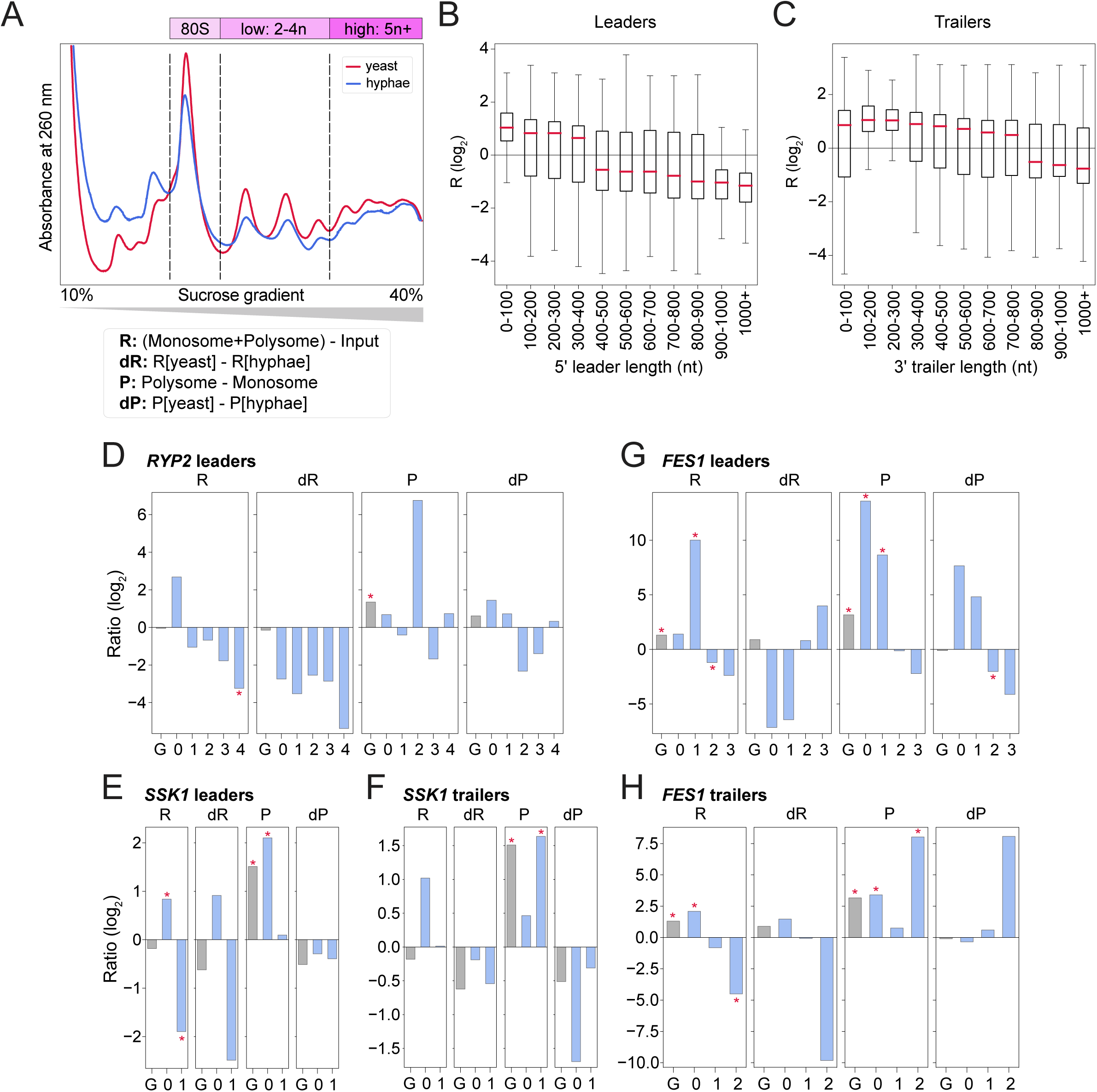
Long 5’ leader and 3’ trailer isoforms correlate with distinct translational states in yeast and hyphae. A) Polysome profiles of yeast and hyphae are shown. Steady-state yeast and hyphal cells were lysed and separated on 10-40% sucrose gradients. Absorbance at 260 was recorded and fractions were pooled into monosome, low polysome, and high polysome for RNA extraction and Illumina RNA-seq. Four metrics from a linear model fit to this data are presented: R (monosome + polysome relative to input), dR (differential R between yeast and hyphae), P (polysome relative to monosome), and dP (differential R between yeast and hyphae). B) Box plots show R values of all isoforms grouped by 5’ leader length bin. Only isoforms of genes encoding both long and short 5’ leader isoforms are included. Red bars indicate the median, and boxes indicate interquartile range (IQR) (Spearman correlation coefficient rho: −0.354, p=1.43e-194; Pearson correlation coefficient r: −0.301, p=1.61e-138). C) Box plots show R values of all isoforms grouped by 3’ trailer length bin. Only isoforms of genes encoding both long and short 3’ trailer isoforms are included. Red bars indicate the median, and boxes indicate interquartile range (IQR) (Spearman correlation coefficient rho: −0.262, p=2.54e-221; pearson correlation coefficient r: −0.219, p=9.08e-153). D-H) R, dR, P, and dP comparisons of *RYP2, SSK1* and *FES1* isoforms grouped by 5’ leader and/or 3’ trailer clusters. G denotes gene-level ratio and is plotted in gray; cluster 0 is the shortest. *’s indicate ratios significantly different with a 1.5-fold change threshold and 5% FDR.

We first determined the relationship between transcript-end length and ribosome association by grouping 5’ leader and 3’ trailer isoforms into 100 nt bins based on length and comparing their *R* values. This analysis was restricted to genes encoding both long and short isoforms (**Fig 6B, 6C)**. Isoforms with 5’ leaders longer than 400 nt and 3’ trailers longer than 800 nt showed reduced ribosome association. In contrast, *P* showed no clear dependence on leader or trailer length **(Fig S6)**. Thus, extreme 5’ leader and 3’ trailer lengths were associated with reduced overall ribosome engagement but not with a systematic redistribution between monosomes and polysomes.

Because uORFs in the long *RYP2* 5’ leader were associated with reduced CDS translation, we examined *R* and *P* across *RYP2* 5’ leader clusters (cluster 0, shortest; cluster 4, longest) and at the gene level (**Fig 6D**). Isoforms from clusters 3 and 4 showed reduced ribosome association, although the reduction was significant for only cluster 4. For *SSK1*, the short 5’ leader isoform (cluster 0) showed significant ribosome and polysome association (positive R and P), whereas the long isoform (cluster 1) was significantly ribosome depleted (negative R) (**Fig 6E**). Conversely, the long 3’ trailer isoform of *SSK1* was significantly enriched in polysome fractions whereas the short trailer isoform was not (**Fig 6F**). These results illustrate that 5’ leader and 3’ trailer variation of a given gene can be associated with different translational states.

*FES1*, which encodes an Hsp70 nucleotide exchange factor, showed similarly complex isoform behavior. Short 5’ leader cluster 1 was significantly enriched in polysomes while longer clusters (2 and 3) were not (**Fig 6G**). The shortest 3’ trailer isoform of *FES1* showed greater ribosome association while the longest isoform cluster did not. Both longest and shortest *FES1* trailer isoforms were significantly enriched in polysomes, although the enrichment was greater for the longest isoform (**Fig 6H**).

Together, these findings show that 5’ leader and 3’ trailer isoform diversity is associated with distinct ribosome and polysome association patterns for genes involved in thermal dimorphism and stress signaling. These associations suggest that transcript 5’ and 3’ end choices contribute to morphology specific translational regulation. Future studies will focus on further dissecting the connection between translation regulation and thermal dimorphism with long and short isoform switching.

## Discussion

Molecular diversity of RNA isoforms is well established in humans and model organisms, yet isoform diversity and its contribution to gene regulation remain underexplored across a wide range of eukaryotes, particularly in fungi. Fungi occupy diverse environmental niches and execute distinct developmental programs, making them powerful systems to study isoform-driven post-transcriptional regulation, where isoform choice determines regulatory capacity. Motivated by *Histoplasma*’s thermal dimorphism, we reasoned that temperature-dependent isoform expression and subsequent post-transcriptional regulation could be central to precise temperature-responsive establishment and maintenance of yeast and hyphal developmental programs. We used an integrated transcriptome annotation framework combining Nanopore direct RNA-seq, Illumina RNA-seq, and 5’-seq, and found that the *Histoplasma* transcriptome harbors extensive isoform diversity through alternative 5’ leaders, 3’ trailers and splicing. Temporal TSS analysis following an acute temperature shift from 37°C to 22°C revealed rapid temperature-responsive shifts in TSS selection, indicating a relatively rapid ability of the cells to alter TSS availability. Some longer 5’ leader isoforms contained regulatory RNA features such as uORFs, and some longer 5’ leader and 3’ trailer isoforms correlated with isoform-specific patterns of ribosome and polysome association. Together, our results indicated that temperature- and morphology-associated isoform diversity had clear consequences for translation regulation. Additionally, the rapid nature of isoform regulation in response to temperature leads to the compelling speculation that it plays an important role in influencing thermal dimorphism and other temperature-dependent biology. Our efforts position *Histoplasma* as an emerging eukaryotic system for studying temperature-responsive RNA isoform biology, in which precise thermal control likely has profound developmental consequences.

RNA isoform diversity can arise from various alternatively regulated or stochastic events during transcription and pre-mRNA processing. Alternative promoters that give rise to new TSSs can create longer 5’ leader isoforms. On the 3’ end, alternatively controlled transcription termination and cleavage and polyadenylation can give rise to diverse 3’ trailer isoforms. Complicating matters, alternative splicing events in coding regions can diversify the proteome and alter protein domain architecture whereas alternative splicing events in noncoding regions can change the 5’ leader and 3’ trailer composition. Our data show that the diversity of RNA isoforms can rapidly respond to external cues, within two hours, with likely regulatory consequences.

We found that *Histoplasma* 5’ leaders and 3’ trailers are substantially longer than those of other fungi (**Fig 2A and 3A**). Notably, *Histoplasma* 3’ trailers were even longer than those of the multicellular metazoans *D. melanogaster* and *C. elegans*. It is unclear whether these noncoding RNA regions are necessary for optimal organismal fitness. Although introns are under selective evolutionary pressure since they determine functionality of proteins, it is harder to extend this logic to 5’ leaders and 3’ trailers since their lengths do not alter the protein sequence but rather protein production and abundance^67^. It is plausible to hypothesize that long 5’ leaders and 3’ trailers would be actively selected for if they include RNA elements or features that drive post-transcriptional molecular fitness in terms of translation and decay^68^. Interestingly, 3’ trailer length has been correlated with morphological complexity in metazoans^69^. This relationship does not appear to generalize to fungi since the 3’ trailers in the multicellular fruiting mushroom *C. cinerea* are much shorter than those in *Histoplasma,* despite its greater morphological complexity. Although the evolutionary basis of longer 5’ leaders and 3’ trailers remains unknown, there is ample evidence that these regions regulate gene expression during stress and development by modulating post-transcriptional processes, consistent with the temperature-responsive isoform expression and translation patterns observed in our study.

We identified many genes with alternative TSSs that produce longer 5’ leader isoforms (**Fig 2**). An outstanding question is which factors control TSS selection across temperatures and morphological transitions in *Histoplasma*. Work in other eukaryotes shows that TSS choice is shaped by interplay among transcription factors and chromatin regulators^70,71^. Use of distal promoters can generate upstream TSSs and longer 5’ leader isoforms while repressing CDS-proximal TSSs through transcriptional interference. In *Saccharomyces*, TSS switching to use more distal promoters, and resultant transcriptional interference, has been linked to chromatin regulators including the histone methyltransferases Set2, where transcription from the distal promoter directs Set1- and Set2-dependent histone methylation which suppresses transcription^72^. Additionally, the chromatin remodeling complex Swi/Snf2 is shown to contribute to LUTI-based interference with the downstream promoters^73^. Consistent with a role for chromatin regulators, recent work from our laboratory identified that the histone deacetylase Rpd3 is required for yeast-phase growth at 37°C and *RPD3* deletion resulted in inappropriate hyphal growth at 37°C^43^. Notably, chromatin immunoprecipitation (ChIP) sequencing detected histone 3 lysine 9 and 14 acetylation (H3K9Ac, H3K14Ac) that were modulated by Rpd3 near the *WET1* distal TSS associated with the longer 5’ leader in yeast cells. This suggests that chromatin regulation by Rpd3, either directly or indirectly, influences accessibility at the *WET1* distal promoter and future work will determine if this observation holds true for other isoforms.

Transcription factors have been shown to orchestrate alternative TSS selection. Although the direct mechanism and key players of alternative TSS selection in *Histoplasma* remain unknown, several patterns suggest plausible mechanisms. Because alternative TSS usage, and therefore expression of longer 5’ leader isoforms, is both temperature-responsive and correlated with distinct developmental states, transcription factors whose functions are temperature or morphology-dependent that are regulators of thermal dimorphism - including Ryp1-4, Wet1, Vea1, and Stu1 – are strong candidates for determining TSS selection during temperature-dependent development. Notably, many of these important transcription factors themselves have morphology-dependent 5’ leader and 3’ trailer isoforms (**Fig 2 and 3**).

In addition to suggesting regulatory mechanisms, our data provide precise transcript annotations, thereby increasing the quantitative power of many gene expression analysis approaches by allowing more accurate read mapping^74,75^. Isoform- and gene-level differential expression analyses of yeast vs hyphae (Y/H) comparison yielded similar expression profiles whether old or new transcript models were utilized, however some genes had clear improvements in quantification. Additionally, we observed that isoform expression levels did not always correlate with overall gene expression. For example, a gene whose overall expression is higher in yeast might encode certain isoforms that are more abundant in hyphae. Globally, we explored the Y/H differential expression of long and short 5’ leader and 3’ trailer clusters to dissect whether both long and short clusters were co-expressed in different morphological states. Differential expression of long and short 5’ leader isoforms was poorly correlated (Pearson’s r: 0.360), suggesting that TSSs that give rise to long or short 5’ leader isoforms could be independently regulated in yeast or hyphae, or that the utilization of distal promoters interferes with the utilization of the proximal promoters.

We identified morphology-specific splice isoforms, some of which altered protein domain lengths and others of which changed leader or trailer architecture. Even though alternative splicing events are widely characterized in multiple eukaryotes, they are not systematically explored in thermally dimorphic fungi. We noted that alternative splicing of the last intron of *RYP2* produced an isoform with an extended C-terminal prion-like domain, potentially influencing localization, phase separation, and intermolecular interactions^76,77^ (**Fig 2**). In addition to impacting protein sequence, we identified many introns in the 5’ leader and 3’ trailer regions which could alter the inclusion of regulatory elements in the RNA.

Notably, diverse 5’ leader, splicing and 3’ trailer isoforms were enriched in many known regulators of thermal dimorphism, such as Ryp2 and Drk1. While longer 5’ leader isoform-encoding genes only account for ∼20% of all protein coding genes, 66.6% (six out of nine members) of the Drk1 hybrid histidine kinase pathway, inferred from orthologs^78,79^, had long and short 5’ leader isoforms with diverse Y/H differential expression trends. This suggests that isoform diversity and downstream post-transcriptional processes likely play regulatory roles throughout establishment and maintenance of thermal dimorphism. Together, these results indicate that longer 5’ leaders are more widespread in *Histoplasma* than previously known (2,328 genes in this study vs. 187 genes reported previously). Morphology-biased long and short 5’ leader isoform expression is enriched among transcription factors and kinases including key regulators of thermal dimorphism. These data link 5’ leader isoforms with unique morphology-specific expression signatures to core transcriptional and signaling regulators of thermal dimorphism. This raises the possibility that alternative 5’ leaders help control and fine-tune thermal dimorphism post-transcriptionally.

We observed that a subset of genes underwent rapid TSS switching upon temperature shift, indicating that isoform switching can be exquisitely temperature responsive. These switches were detected 2 hours post-temperature shift even though the *Histoplasma* doubling time is greater than 9 hours and the morphological transition is not apparent until days 4-10. Other genes did not undergo TSS switching until the morphologic transition was established, indicating differential regulation of isoform production that could relate to biological functions of genes in development.

An important question raised by long and short 5’ leader isoform switching is which cis-regulatory elements in longer leaders confer functional differences such as translation and RNA decay. Our results point to uORFs as one such element. This focus on uORFs is supported by a systematic analysis of cis-acting 5’ leader elements in yeast which shows that uAUGs and uORFs are among the strongest determinants of gene expression, and by prior work linking LUTIs to uORF-mediated translational control during meiosis in *S. cerevisiae*^20,26^. Although our analysis likely captured only a small subset of uORFs since we required a minimum ORF length and ribosome footprint evidence to designate a uORF as translated, we nonetheless identified a small group of genes in which uORF presence in the longer 5’ leader isoform is associated with isoform-specific translational repression in either yeast or hyphae. The modest number of genes with uORFs suggests that translation regulation due to long and short 5’ leader isoform switching is likely not universally mediated by uORFs and likely mediated by multiple cis-regulatory elements and post-transcriptional mechanisms. Detecting more uORFs may require relaxing our criteria of uORF length and ribosome footprint signal. Furthermore we could account for rapid decay of uORF-containing longer 5’ leader isoforms through likely nonsense-mediated decay^9,19^. Collectively, these findings indicate that isoform switching can expose or hide translational control elements in 5’ leaders. This motivates future searches for additional cis-regulatory features, beyond uORFs, that could regulate gene regulation throughout acute temperature changes and thermal dimorphism.

To evaluate translation regulation at the isoform-level we used polysome-seq to determine the distribution of long and short isoforms into monosome-associated and polysome-associated populations. Despite baseline differences in the polysome profiles that we obtained from yeast and hyphae, it was clear that some long and short isoforms had distinct translational states. Consistent with our uORF analysis and uORF-mediated translational repression, the longer *RYP2* 5’ leader cluster containing the uORF (cluster 4) had the lowest ribosome association among other *RYP2* 5’ leader clusters. We are particularly intrigued by the hypothesis that longer leader isoforms may play a role in the transition between temperatures: for example, the regulatory elements in some longer leaders might be refractory to translation at one temperature (e.g. 37°C) but permissive for translation at another temperature (e.g. 22°C). Such a mechanism might allow the initial production of hyphal-specific proteins by yeast cells that are shifted to 22°C until a shorter leader transcript is produced and translated. Future work will include mechanistic dissection of *Histoplasma* 5’ leaders and 3’ trailers, and their biological consequences using reporter systems and targeted perturbation of specific isoforms, ultimately exploring how RNA isoform diversity determines the post-transcriptional principles underlying thermal dimorphism.

## Materials and Methods

### Experimental Methods

#### *Histoplasma* growth and maintenance

*Histoplasma ohiense* strain G217B obtained from the American Type Culture Collection (ATCC, 26032) was used for all experiments. Yeast cells were thawed from frozen stocks and grown on *Histoplasma* Macrophage Medium (HMM) agarose plates at 37°C with 5% CO_2_. Both yeast and hyphal cells were grown in liquid HMM media without glucose supplemented with 100 mM N-acetyl-glucosamine (GlcNAc), hereafter HMM+GlcNAc, on an orbital shaker at 120 rpm. Yeast liquid cultures were grown at 37°C with 5% CO_2_ and hyphal cells were cultured at 22°C without CO_2_. Yeast cells were cultured in a Biosafety Level 2+ (BSL-2+) facility and hyphal cells were cultured in a Biosafety Level 3 (BSL-3) facility.

#### Culture conditions for steady-state yeast and hyphal growth

Yeast cells grown on HMM agarose plates were used to inoculate 5 mL HMM+GlcNAc media. The 5 mL culture was grown for three days, passaged 1:25, and grown for 2 days. To establish a steady-state yeast culture, yeast cells from the 2-day culture were used to inoculate a larger culture where the optical density at 600 nm (OD_600_) would be at 5-8 the next day. To establish a steady-state hyphal culture, yeast cells from the 2-day culture were used to inoculate a larger culture at starting OD_600_ = 0.025. This low-density culture was grown for 7-10 days and then passaged twice at 1:100, every 7 days, to maintain a steady-state hyphal culture. Yeast cells were pelleted by centrifugation and hyphal cells were collected by filtration through a 0.45 µm filter. 1 mL QIAzol (or TRIzol) was added to each pellet, cells were flash frozen in liquid nitrogen and stored at −80°C.

#### Culture conditions for 37°C to 22°C transition

Mid-log yeast cultures grown at 37°C were transitioned to a 22°C water bath with an orbital shaker for 30 minutes to ensure quick adjustment of temperature. Flasks were then placed in the dry shaking incubator at 22°C and 120 rpm. Cells were pelleted after 2 hours, 6 hours, 24 hours (day 1), and 96 hours (day 4) post-temperature shift. 1 mL QIAzol (or TRIzol) was added to each pellet after which pellets were flash frozen in liquid nitrogen and stored at −80°C.

#### RNA extraction

Cell pellets in QIAzol were thawed and transferred to 2-mL tubes with 0.5 mm diameter zirconia beads. Pellets were subjected to bead beating in Trizol for 2 minutes, 200 µL chloroform was added to each tube followed by vortexing for 30 seconds. Tubes were incubated at room temperature for 2 minutes and centrifuged at 21,000 x g for 15 minutes. The aqueous layer was transferred to a new tube and an equal volume of 70% ethanol was added. The resulting aqueous phase was transferred to EconoSpin RNA columns (Epoch Life Sci 1940-250). To extract RNA, columns were subjected to a wash with Buffer 1 (900 mM guanidium thiocyanate, 20% ethanol, 10 mM Tris-HCl pH 7.5). RNA was subjected to on-column DNase treatment using the Qiagen RNase-free DNase kit. Then, columns were subjected to two sequential washes with Buffer 1 and two sequential washes with Buffer 2 (100 mM NaCl, 80% ethanol, 10 mM Tris-HCl pH 7.5) prior to elution with nuclease-free water. RNA quality was evaluated by using either an Agilent Bioanalyzer with an RNA 6000 Nano chip, Tapestation with RNA ScreenTape, or an agarose gel.

#### Nanopore direct RNA-seq and Illumina RNA-seq

RNA from steady state yeast and hyphal cells were used as input to isolate mRNA by using the NEBNext High Input poly(A) mRNA isolation module. Five independent biological replicates were analyzed. 5’ biotinylated RNA oligo was ligated to select full-length capped RNA molecules as described previously^80^. Briefly, 3 µg of poly(A) mRNA was dephosphorylated using 75 units of Calf Intestinal Alkaline Phosphatase at 37°C for 1 hour. Dephosphorylated mRNA was purified using Phenol:Chloroform:Isoamyl Alcohol (25:24:1 – hereafter PCI) and precipitated using 1-volume 5 M ammonium acetate pH 5.2, 2.5-volume ethanol and 1 µL of GlycoBlue. Dephosphorylated RNA was then decapped using 100 units of mRNA decapping enzyme at 37°C for 2 hours. RNA was purified using PCI and precipitated as described. Decapped RNA was then used to ligate a 5’ biotinylated oligonucleotide (ssRNA Oligo: 5’-biotin-CGACUGGAGCACGAGGACACUGACAUGGACUGAAGGAGUAGAAA-3’, IDT). Ligation was conducted using 0.25 µg of oligonucleotide in the presence of T4 ligase 1, ATP and PEG8000 at 16°C for 16 hours. RNA was purified using PCI and precipitated as described. mRNA with ligated oligonucleotide was pulled down using Streptavidin MagneSphere Paramagnetic Particles (Promega, Z5481). Enriched full-length mRNA with 5’ biotinylated oligonucleotide was subjected to Nanopore Direct RNA Sequencing through SeqCenter (Pittsburgh, PA). Libraries were prepared using the PCR-free Oxford Nanopore Technologies (ONT) Ligation Sequencing Kit (SQK-RNA004) with the NEBNext Companion Module (E7180L) according to manufacturer’s instructions.

Illumina sequencing libraries were prepared using the matching total RNA samples from replicates 1-3 from the Nanopore direct RNA-seq experiment. Library preparation was performed at SeqCenter using Illumina stranded mRNA prep and 10 bp unique dual indices (Illumina #20040534). Sequencing was done on a NovaSeq X plus platform, and produced paired-end 150 bp reads.

#### 5’-seq (STRIPE-seq)

RNA from steady-state yeast or hyphal cells, or from 37°C to 22°C transition, was used as input for 5’-seq library preparation. 5’-seq libraries were prepared using the previously published STRIPE-seq protocol (here referred to as 5’-seq for simplicity)^81,82^. All oligonucleotide sequences can be found in the Supplemental Table S5 from Policastro et al., 2020^81^. Briefly, 50-200 ng total RNA samples were treated with 0.2 units of Terminator 5’-Phosphate-dependent Exonuclease (TEX) in a 2 µL reaction to preferentially degrade uncapped RNAs at 30°C for one hour. 2 µL decapping reaction was then used for reverse transcription and template-switching reactions. 2 µL of the decapping reaction was mixed with the master mix including 1.5 µL of sorbitol/trehalose solution (3.3 M sorbitol and 0.66 M trehalose), 1 µL RTO (reverse transcription oligo, 10 µM), 0.5 µL dNTPs (10 mM). Then, the mixture was incubated at 65°C for 5 min, and 4°C for 2 min to anneal RTO to RNA. A master mix containing 2 µL betaine (5 M), 2 µL SuperScript II First Strand Buffer, 0.5 µL DTT (0.1 M), and 0.5 µL SuperScript II Reverse Transcriptase was added to the annealed samples. The reverse transcription reaction (with the final volume of 10 µL) was incubated at 25°C for 10 min followed by 42°C for 5 min. Immediately after, 0.25 µL of TSO (template-switching oligo, 400 µM) was added, and incubated at 42°C for 25 min followed by 70°C for 10 min. The products were cleaned up using 8 µL of SPRI beads for the 10 µL reaction, and beads were eluted using 12 µL nuclease-free water and 11 µL were transferred to a new tube. Libraries were prepared using 0.75 µL of FLO (forward library oligo, 10 µM), 0.75 µL RLO (reverse library oligo, 10 µM) and 12.5 µL 2x KAPA HiFi HotStart Ready Mix along with the 11 µL cleaned template-switching product. PCR products were then size selected by SPRI double size selection. Final libraries were evaluated on Agilent bioanalyzer or tapestation prior to sequencing. Libraries were sequenced at UCSF CAT (Center for Advanced Technology) on a NovaSeq X plus platform, and produced paired-end 150 bp reads.

#### Polysome profiling, fractionation, and sequencing

Steady-state yeast and hyphal cells were grown as described above. Cells were treated with 100 µg/mL cycloheximide for 2 minutes and pelleted. Cell pellets were flash frozen in liquid nitrogen and stored at −80°C. Pellets were mixed with 1 mL frozen polysome lysis buffer (PLB: 20 mM Tris pH 8, 140 mM KCl, 5 mM MgCl2, 0.5% Triton X-100, 0.5 mg/mL heparin, 200 µg/mL cycloheximide, 1 mM DTT, 200 U SUPERase.In RNase inhibitor). Frozen pellets along with PLB were subjected to three two-minute rounds of cryogenic grinding at 30 mHz (Mixer Mill MM 400, Retsch). The resulting cell powder was transferred to 50 mL conical tubes and stored at −80°C. Cell powders were thawed on ice by adding 1 mL ice-cold PLB on ice. Then lysates were clarified by centrifuging at 20,000 x g for 2 min at 4°C. Clarified lysates were passed through 0.2 µm steriflip filters twice to remove any remaining infectious particles and safely removed from the BSL-3 facility. RNA concentration of the lysates was determined by Nanodrop, and 200 µg equivalent of cell lysates were layered on 10-40% sucrose gradients. Briefly, sucrose gradients were prepared as follows. 10% of 40% sucrose solutions (10% or 50% sucrose, 20 mM Tris pH 8, 140 mM KCl, 5 mM MgCl2, 0.5 mg/mL heparin, 100 µg/mL cycloheximide, 20 U/mL SUPERase.In RNase inhibitor) were layered in 13.2 mL thin wall polypropylene tubes (Beckman, #331372). Sucrose gradients were prepared using Gradient Master 108 (Biocomp). Loaded gradients were centrifuged at 39,000 x g for 2 hours at 4°C using Beckman L8-80M ultracentrifuge and SW41-Ti rotor. 10% of the lysates were saved as input. Centrifuged sucrose gradients were fractionated into 400 µL fractions using piston gradient fractionator (Biocomp), and absorbance was recorded at 260 nm to evaluate polysome traces. Fractions that correspond to monosomes, low polysomes (2-4n), and high polysomes (5n+) were pooled with a final volume of 400 µL per pool. RNA from each pool was extracted using 800 µL TRIzol and 300 µL 24:1 chloroform:isoamyl alcohol. Aqueous phase was then cleaned up using Zymo RNA clean and concentrator kit. Purified RNA was then precipitated using equal volume 7.5 M LiCl to remove heparin. RNA quality was evaluated using Nanodrop and Agilent tapestation. RNA samples were then subjected to rRNA depletion prior to library preparation and sequencing. To deplete rRNA, we designed custom oligos using Oligo-ASST against *Histoplasma* rRNA sequences as predicted by RNAmmer v1.2 in eukaryote mode **(Table S1)**. Oligo pool with 257 oligos was synthesized at Integrated DNA Technologies with expected concentration of 50 pmol per oligo. Up to 2 µg of RNA (in 6 µL) was incubated with 1 µL of oligo pool (final concentration of 400 nM per oligo), 2 µL 5x hybridization buffer (1 M NaCl and 500 mM Tris pH 7.5), and 1 µL 0.1 mM DTT. Hybridization reaction was incubated at 95°C for 2 min, and then gradually cooled to 22°C (0.1°C/second). Hybridization reaction was treated with 10 U of RNase H to degrade rRNA, followed by DNase treatment to degrade oligos. rRNA-depleted RNA samples were then cleaned up using Zymo clean and concentrator kit. RNA samples were sequenced at SeqCenter using Illumina stranded total RNA prep and 10 bp unique dual indices (Illumina #20040534). Sequencing was done on a NovaSeq X plus platform, and produced paired-end 150 bp reads.

## Data Analysis Methods

### Sources of *Histoplasma* genome assembly and annotation

The UCSF3 assembly of *Histoplasma ohiense* G217B (GCA_051313235.1) was used as genomic reference^83^. The corresponding gene annotation, which is a BLAT-based lift of the Illumina RNA-seq based annotation of Gilmore *et al* 2015^49^, was used as the reference transcriptome, and the corresponding LTRHarvest^84^ and TBLASTN^85^ based transposon annotations were used for transposon filtering.

### Nanopore direct RNA-seq data analysis

Nanopore direct RNA-seq reads were filtered for full-length reads by using BLASTN^85^ to locate a ligated 5’-adapter. Reads with BLASTN hits with a maximum e-value of 100 and coverage of at least 10 nucleotides of the ligated adapter were included as full-length. Filtered reads were then mapped using minimap2^86^ v2.24 (-ax splice -k 14 -uf -G 2000). Both 5’-adapter selected and all reads were subjected to further quality filtering by removing unmapped reads, reads with split alignment BAM tags and reads that were smaller than 200 bp.

### 5’-seq data analysis

5’-seq reads were analyzed using TSRexploreR^87^ (git commit 0b6cca0b42) to determine TSSs at the single nucleotide resolution. Prior to TSRexploreR, paired-end reads were cleaned by removing rRNA reads using HISAT2^88^ v2.2.1. Then, unique molecular identifiers (UMIs) were extracted and included in the headers using *umi-tools extract*^89^. Cutadapt^90^ v4.2 was then utilized to enforce the correct R1 read structure that included “TATAGGG”, which is the expected sequence based on the presence of template-switching oligo. Another pass of cutadapt was run to remove template switching chaining. Final filtered reads were then mapped to the reference UCSF3 genome using HISAT2. Then, mapped reads were deduplicated using *umi-tools dedup*. Secondary alignments and reads without mates were also removed from the final BAM file. Final BAM files were then used as inputs for TSRexploreR and TSS sites were determined and exported as bedgraph files.

### Genome-guided transcriptome assembly

FLAIR3 (full-length alternative isoform analysis of RNA v3) was used to annotate transcript models using long-read, short-read, and 5-seq data. All yeast and hyphal reads from either 5’-adapter filtered or non-filtered files were merged. We ran FLAIR using all four merged read files as input for the FLAIR align-correct-collapse-combine pipeline along with splice junctions that were extracted from STAR^91^ v2.7.10 alignment of matching Illumina RNA-seq reads and TSS coordinates that were extracted from TSRexploreR that were padded by 50 bp from both 5’ and 3’ directions. Independently, we used the flair transcriptome module using the same input read files. By using both pipelines, along with 5’-adapter filtered and non-filtered reads, we built the first version of the transcriptome assembly by combining (combine parameters: -w 25 -f 5 -p 5) the transcript models generated by both collapse and transcriptome modules. All FLAIR invocations are included in the supplementary materials. This combinatorial approach was chosen to increase our ability to identify 5’ leader and 3’ trailer isoforms, and we then used our internally developed pipeline to validate transcript models identified by FLAIR.

For each FLAIR isoform, a coding sequence (CDS) was annotated as the largest open reading frame (ORF) in the spliced sequence. CDS were annotated only if they began with ATG, ended with a stop codon, and were at least 180 bp.

### 5’ validation with 5’-seq

For 5’ validation, we considered FLAIR isoforms and TSRexploreR TSSs that did not overlap annotated transposons, tRNAs, or rRNAs. TSSs were further required to be supported by at least 10 reads. These TSSs were combined into transcriptional start regions (TSRs) by merging TSSs within 25bp of each other. TSRs shorter than 10bp were 3’ padded to a final length of 10bp. FLAIR isoforms were annotated as 5’ validated if their 5’ start overlapped at least one TSR on the same strand **(Table S2)**.

### Mapping to reference transcriptome

To map FLAIR isoforms to the reference transcriptome, we considered FLAIR isoforms and reference genes that did not overlap annotated transposons, tRNAs, or rRNAs. This gave 70,610 FLAIR isoforms and 11,838 reference transcripts which were placed into clusters of transcripts overlapping by at least 1bp on the same strand, giving 17,168 total clusters. For the 7,738 that contained at least one FLAIR isoform and a single reference transcript, the isoforms were assigned to this uniquely mapped reference transcript. 1,427 reference transcripts did not cluster with any FLAIR isoform, and there were 3,566 clusters of FLAIR isoforms with no reference transcript, which were designated Novel isoforms (N).

The remaining 1,238 clusters with at least one FLAIR isoform and more than one reference transcript were reclustered, this time placing transcripts into the same cluster only if their CDS overlapped by at least 1bp on the same strand. This gave an additional 1,895 clusters of FLAIR isoforms uniquely mapped to a reference transcript, 565 reference transcripts not clustering with any FLAIR isoform, and 1,836 clusters of FLAIR isoforms with no reference transcript, which were added to the list of Novel isoforms. The remaining 101 clusters containing at least one FLAIR isoform, and more than one reference transcript were interpreted as single genes incorrectly split into non-overlapping transcripts in the reference.

### Filtering for complete and abundant isoforms

FLAIR isoforms that were not 5’ validated in the 5’-seq analysis were removed. Where this resulted in reference transcripts with no assigned FLAIR isoforms, those reference transcripts were reclassified as Reference-only (RO) isoforms. Then, unfiltered Nanopore reads were quantified against the remaining, 5’ validated, FLAIR isoforms using FLAIR *quant*. FLAIR isoforms with fewer than 5 counts based on FLAIR quant were removed, and reference transcripts were again reclassified as RO where the count filtering resulted in no assigned FLAIR isoforms.

### Resolving fused genes

FLAIR isoforms coding for disjoint proteins were resolved by the following protocol. For each set of FLAIR isoforms assigned to a single reference transcript, the reference transcript and FLAIR isoforms with CDS at least 270 bp were clustered based on CDS overlap of at least 1 bp. Clusters were merged if they contained FLAIR isoforms with 5’ starts within 25bp of each other. Remaining FLAIR isoforms that did not pass the above CDS >= 270 bp filter were assigned to the new clusters based on greatest overlap of genomic coordinates or minimum distance for cases of no overlap. Resulting clusters not including the original reference transcript were assigned as new *Novel* isoforms and the original *Reference FLAIR isoforms* were reannotated with just the FLAIR isoforms assigned by the new clustering. In cases where no FLAIR isoforms were assigned to the reference transcript in the new clustering, that transcript was assumed to be a false positive in the original annotation and removed.

The final 14,210 isoform clusters are referred to as the assembled gene set for the remaining analysis and consist of 1,992 reference transcripts with no assigned FLAIR isoform (Reference-only isoforms), 2,548 clusters of FLAIR isoforms with no assigned reference transcript (Novel isoforms), and 9,670 clusters of FLAIR isoforms assigned to a reference transcript (Reference FLAIR isoforms). This gene set is provided in the supplemental data in GTF **(Table S3)** and FASTA **(Data S1)** formats.

### 3’ validation by DORADO analysis of Nanopore reads

Poly(A) tail lengths and polyadenylation sites were determined using the *--estimate-poly-a* flag while basecalling using DORADO. BAM files with poly(A) tags (pt: tail length, pa: signal coordinates, and mv: move table) were utilized to convert poly(A) sites to genomic coordinates to determine polyadenylation sites (PAS). For 3’ validation, polyA sites were combined into polyadenylation clusters (PACs) by merging sites within 25 bp of each other. FLAIR isoforms were annotated as 3’ validated if their 3’ stop was within 25 bp of at least one PAC on the same strand.

### Gene and isoform annotation

For reference mapped genes, short names, annotations, repeat context, and ortholog mappings were propagated from the original annotation of Inglis et al. 2013^92^ as updated in Gilmore et al. 2015^49^, Voorhies et al. 2022^93^, and Heater et al. 2026^83^. Transcription factor (TF) annotations were taken from the joint TF annotation of *Histoplasma ohiense* G217B and *Coccidioides posadasii* Silveira described in Homer et al. 2025^94^. Additionally, protein sequences were translated from each coding isoform in the new assembly and Pfam domains in these sequences were annotated with InterProScan v107.0. Full gene and isoform level annotations are given in **tables S4 and S5** respectively. Additionally, protein localization of translated isoform sequences was predicted with DeepLoc2^95^ in Accurate mode **(Table S6)**.

### BLAST analysis of novel gene models

In order to compare the coding *Novel isoforms* set to gene annotations other than the reference annotation, BLASTP from NCBI BLAST+ 2.12.0^96,97^ was used to search the longest translated protein sequence for each *Novel isoforms* gene against the annotated protein sequences of 41 fungal genomes (as listed in Table S18 of Gilmore et al 2015^49^). Results of this analysis are provided in **Table S7**.

### Differential splicing analysis

Paired-end Illumina reads were aligned to the reference genome using HISAT2. The resulting per-replicate BAM alignment files were used as inputs to RMATS_TURBO^98^ v4.3.0 invoked on the assembled gene set GTF with --readLength 150. Differential splicing events were considered significant if they had an adjusted p-value less than 5% and delta psi of at least 10% **(Table S8)**.

### Differential expression analysis

Transcript abundances were quantified from the paired-end Illumina reads for the individual isoforms of the assembled gene set **(Data S1)**. Estimated counts (est_counts) of each transcript in each sample were generated by using kallisto^74^ v0.48.0. Kallisto estimated counts for each isoform in each sample were depth normalized to log2(cpm) values, and between sample normalization was carried out using trimmed median of means (TMM) normalization as implemented in the calcNormFactors function of edgeR. Differentially expressed isoforms were identified by comparing replicates for yeast and hyphal samples using the glmQLFit, glmQLFTest, and topTags functions in edgeR^99^ v4.0.16. Isoforms were considered significantly differentially expressed if they were statistically significant (at 5% FDR) with an effect size of at least 2x (absolute log2 fold change ≥ 1) for a given contrast. Differentially expressed genes were identified in the same way but summing counts over all isoforms **(Table S9)** of a given gene prior to running edgeR. The results of both edgeR analyses are merged in **Table S10**.

### Clustering isoforms by 5’ leader or 3’ trailer

Isoforms were combined into clusters if their 5’ starts were within 50 bp of each other. For each cluster, kallisto estimated counts were summed over all isoforms in the cluster for each sample. Clusters with fewer than 10 summed counts in any sample were removed. Clusters passing this count filter were added to the final set of “multileader clusters” if there were at least two such clusters for a given gene. Additionally, the multileader clusters were reduced to a “short/long leader” set by taking the cluster with the 3’-most start for each gene as the “short” cluster and combining the remaining clusters (and likewise summing their counts for each sample) to give a “long” cluster. “Multitrailer clusters” and “short/long trailer” clusters were defined by the same protocol, except that clustering was on 3’ stops rather than 5’ starts and the “short” cluster was taken as the cluster with the 5’-most stop. Genes in the multileader and multitrailer clusters were subjected to gene ontology analysis using InterPro2GO mapping along with manual curation.

### Differential expression of isoform clusters

For each of the four defined sets of isoform clusters; viz., multileader, multitrailer, short/long leader, and short/long trailer; differentially expressed genes were identified using edgeR, running separate analyses for each of the four sets. In each analysis, in addition to the summed counts for each cluster, gene level summed counts were included for all genes that did not have multiple clusters in that analysis. In this way, the same number of total counts over isoforms were included in each analysis. This was necessary in order to generate an equivalent cpm depth normalization in each analysis and to supply approximately equivalent variance distributions for the empirical Bayes step used for calculating p-values in edgeR. Otherwise, the same protocol as was used for the gene-level analysis was used for depth and TMM normalizations, dispersion estimation, and fitting of the clustered isoform sets. The results of these analyses, and the subsets identified by the above gene ontology analyses, are provided in **tables S11-S20**.

### Principal Components Analysis

Principal components analysis (PCA) was applied at several points in our analysis. In all cases, PCA was implemented by using the svd function from the linalg module of numpy (version 1.24.2)^100^, giving the principal components as left singular vectors. Variances of each component were calculated as the squares of the singular values and these variances were normalized by dividing by total variance to give relative variances. Numerical rank was defined as the number of components with relative variance at least 20%. Unless otherwise noted, data matrix columns were mean centered prior to SVD.

### Reclustering TSSs to TSRs for time course analysis

Our method of aggregating TSSs to TSRs described above for 5’ validation is optimized for sensitivity at the cost of spatial and temporal specificity. For time course analysis of transcriptional start sites, we inverted this sensitivity/specificity trade off by repartitioning the TSSs into narrower TSRs as follows. First, TSRexploreR was re-run as before, except that separate TSS sets were generated for each timepoint and filtered for at least 10 counts over all replicates of that timepoint. These state specific TSSs were then mapped to the original set of TSRs that were generated from the pool of all samples. For each original TSR, the corresponding TSSs were aggregated on overlap to generate a consistent set of TSS coordinates over all timepoints. TSRs for which no timepoint-specific TSSs were mapped, due to the increased stringency of the count filter, were removed from the time course analysis. To identify contiguous sequences of covarying TSSs, we generated a matrix of counts for each aggregated TSS at each timepoint for each TSR and performed PCA on this matrix without centering. Peaks were defined as the set of unique genome locations corresponding to the absolute maximum weights of each PC with relative variance >= 20%, and each TSS was assigned to the closest peak. For each peak, assigned TSSs within 25bp of each other were reclustered to generate a new set of TSRs. The net effect of this procedure was to split the original TSRs at locations of divergent expression and to slightly shrink the TSRs due to more stringent count filtering at the level of individual timepoints. Final reclustered TSR coordinates are provided in **Table S21**.

### Differential expression of TSRs

TSSs were detected and quantified from genome aligned 5’-seq reads by TSRexploreR independently for each sample in the steady state yeast and hyphal cultures and in the 37°C to 22°C transition. The TSS quantification was then aggregated to the reclustered TSRs by summing counts for all TSSs overlapping a given TSR, and analysis was restricted to TSRs with at least 10 counts in at least 6 samples. To further restrict the analysis to high confidence TSRs, we applied an additional filter based on the relative TSR abundance for each gene. Specifically, we calculated depth normalized TSR abundance as log2 counts per million with a pseudocount of 1, *i.e.* log_2_(cpm). Then for each gene, we calculated the maximum abundance of each TSR as its maximum log_2_(cpm) over all samples and removed any TSR with maximum abundance less than 10% of the highest maximum abundance for that gene.

Summed TSR counts were used as input to edgeR to detect significant differential expression between steady state yeast and hyphae or between the initial (t=0) and subsequent timepoints in the 37°C to 22°C transition. TSRs were considered differentially expressed if they had at least a 2-fold change at FDR < 0.05 for the steady state Y/H contrast or at least a 1.5-fold change at FDR < 0.05 for at least one time course contrast. The edgeR estimated contrasts for both experiments are given in **Table S22**. Additionally, gene-level expression was estimated by summing the counts for all TSRs of each gene, and differentially expressed genes were defined based on the same edgeR protocol used in the TSR-level analysis.

Genes with diverse TSR expression were detected using gene-level principal component analysis (PCA) as follows. For each gene with at least two significantly differential TSRs, we performed PCA on the matrix of all edgeR estimated contrasts (columns) for all significantly differential TSRs of that gene (rows). In practice, only one gene had numerical rank greater than 2. All genes with numerical rank >= 2 were considered to have diverse expression, and defined as the class 2 gene set. Additionally, genes with numerical rank 1 (*i.e.*, genes for which at least 80% of the variance in TSR contrasts corresponded to a single component) were considered to have diverse expression if there was at least one TSR with positive weight and at least one TSR with negative weight on the first principal component, and were defined by the class 1 gene set. Genes in class 1 and class 2 were further classified based on the 5’→3’ ordering of their expression patterns. For genes in class 1, the projection of the gene level expression pattern on the first component was compared to the projection of each TSR level expression pattern on that component, with TSRs being considered “major” if they had the same orientation as the gene and “minor” if they had opposite orientation. Genes were classified as “dominant short” if all minor TSRs were 5’ of all major TSRs, “dominant long” if all minor TSRs were 3’ of all major TSRs, and “other” for all other patterns. For genes in class 2, we carried out the same classification independently on the first two components, adding a “neutral” category for all major TSRs on a given component (an outcome not possible for class 1 cases). We then classified genes on the pair of individual component classifications (*e.g.*, PC1: neutral, PC2: dominant short). Examples for some of these classifications are shown in the Fig S7. For the final clustered heatmaps, genes were first divided on class, then on 5’→3’ ordering, then on yeast, hyphal, or no enrichment in the steady state comparison (based on 1.1x change with no significance threshold), then on positive (hyphal) or negative (yeast) slope of a line fit to the time course data. The results of the PCA based analysis are provided in **Fig 4B** and **Table S23** (class 1) and **Fig S5** and **Table S24** (class 2).

### Definition of isoform level uORFs

For each assembled gene mapped to a protein coding reference gene, upstream open reading frames (uORFs) were defined over the set of assembled isoforms as follows. Canonical and alternative start codons were taken as AUG, CUG, GUG, UUG, ACG, AUU, AUA, and AUC. Isoforms were spliced to give inferred mRNA sequences. The same intron splicing was applied to transform the annotated start codon of the reference gene onto each inferred isoform mRNA sequence, discarding isoforms that were not annotated with a CDS or that did not transcribe this annotated reference start codon. For the remaining isoforms, uORFs were defined as mRNA subsequences of at least 18 bases (6 codons), starting with one of the above start codons, uninterrupted by an in-frame stop, and terminated by a stop codon 5’ relative to the reference start codon. For uORFs overlapping in the same reading frame, only the largest uORF was included. uORFs identified for different isoforms were considered identical, and analyzed as single entities, if they were transcribed from the same genomic location. uORF genomic coordinates are provided in **Table S25**.

### Re-analysis of ribosome footprints relative to isoform level uORFs

The strand-specific, single-end reads from the ribosome footprint and matched mRNA samples of Gilmore et al 2015^49^ [GEO: GSE68705, SRA: PRJNA283425] were linker-stripped and filtered for mitochondrial, transposon, and rDNA sequences as previously described^49^. The remaining, linker-stripped sequences were mapped to the G217B UCSF3 reference genome with Bowtie^101^ v1.3.0, restricting output to unique alignments (-k1) and post-filtering for full length alignments of the query sequences. Length filters for aligned sequences (22-32bp for ribosome footprint samples and >= 22bp for mRNA samples), correction for first position T artifact, inference of ribosomal P-site were as previously described^49^. FPKMs were then calculated for the aligned reads of each sample type for each reference CDS and each unique isoform-level uORF as previously described^49^ **(Table S26)**.

Translational efficiency (TE) in yeast or hyphae was calculated for uORF or CDS features as ribosome footprint FPKMs over mRNA FPKMs, restricting this calculation to CDS with at least 128 total counts and uORFs with at least 16 total counts in a given morphological state. For both uORFs and CDS in both morphologies, the median TE was near zero. In order to identify candidate genes with morphology-specific uORF-mediated translational repression, we filtered for genes where the CDS had translational repression in one morphology and at least one uORF had ribosome enrichment in the other morphology, where CDS were considered to have morphology-specific translational repression, and uORFs to have morphology specific ribosome enrichment, if there was at least a 2 fold difference between yeast and hyphae or if only one morphology had sufficient counts to calculate TE. We then used the isoform-level kallisto estimated counts (described above) to test for differential expression of isoforms consistent with uORF-mediated translational repression as follows. For each candidate uORF, the isoforms of each candidate gene were split into three clusters: A) isoforms with reference CDS and lacking the uORF, B) isoforms containing the uORF, and C) isoforms lacking both the reference CDS and the uORF; cluster C was discarded, counts were summed independently for clusters A and B, and cases where either clusters A or B had fewer than 10 counts were discarded. This left 93 uORFs on 47 yeast repression candidate genes and 213 uORFs on 83 hyphal repression candidate genes with sufficient counts. Summed counts for A or B clusters were imported into edgeR, along with summed gene level counts for the remaining genes (for cpm and variance estimation consistent with the edgeR protocols above) and clusters with 2-fold differential expression between yeast and hyphae at 5% FDR were identified as above. This identified final sets of 49 uORFs on 31 hyphal repression candidate genes with hyphal enriched uORF containing isoforms (“B” clusters) or yeast enriched isoforms with CDS but not the uORF (“A” clusters) **(Table S27)** and 58 uORFs on 24 yeast repression candidate genes with yeast enriched “B” clusters or hyphal enriched “A” clusters **(Table S28)**.

### Estimating differential expression and differential fractionation from polysome profiling

Due to the fractionation, purification, and sequencing protocols, the ratio of in vivo RNA concentration to quantity of sequenced RNA differed among sample types by unknown scale factors. Therefore, the following analysis focuses on relative changes among each sample type: e.g., rather than directly estimating the ratio of ribosome associated to input RNA for a given isoform, we estimate the difference in that ratio for a given isoform relative to the mean for all isoforms.

To this end, kallisto estimated counts for each isoform in each sample **(Table S29)** were depth normalized to log_2_(cpm) values, and between sample normalization was carried out using TMM normalization as implemented in the calcNormFactors function of edgeR to correct for sample-specific highly abundant isoforms. TMM normalized log_2_(cpm) values were fit to a linear model in edgeR, first estimating dispersions with estimateDisp and then fitting the model with glmQLFit. Fit parameters and adjusted p-values for non-zero parameters were extracted with topTags. The model was y = u + D + R + dR + P + dP + H + dH + epsilon where, for a given isoform, y is observed log_2_(cpm), epsilon is the residual due to counting, biological, and technical variation (assumed to be distributed as a negative binomial); u is the mean log_2_(cpm); D is the difference between yeast and hyphae; R is the difference between ribosome fractions and input RNA; P is the difference between polysome and monosome fractions; H is the difference between high and low polysome fractions; and dR, dP, and dH are the differences between yeast and hyphae for R, P, and H respectively. The model was parameterized so that, for all terms other than u, 0 corresponds to no change, positive fit parameters correspond to higher counts in the numerator (e.g., yeast for the D term), and negative fit parameters correspond to higher counts in the numerator (e.g., hyphae for the D term). For this reason, fit parameter values correspond to half of the log_2_ fold change between the numerator and denominator. The fit values were therefore scaled by a factor of 2 when filtering for differentially expressed genes and these scaled parameter values, which can be interpreted as fold changes, are reported in the tables and figures of this manuscript.

For gene level analysis, isoform level counts were summed for each gene in each sample and then fit exactly as above **(Table S30)**. For analysis of isoforms clustered by leader or trailer **(Tables S31 and S32)**, isoform level counts were summed for each cluster for each gene. Additionally, gene level summed counts were included for all genes that did not have multiple clusters in each analysis. In this way, the same number of total counts over isoforms were included in each analysis. This was necessary in order to generate an equivalent cpm depth normalization in each analysis and to supply approximately equivalent variance distributions for the empirical Bayes step used for calculating p-values in edgeR.

### Defining genes and isoform clusters with morphology dependent polysome vs. monosome fractionation

The dP parameter estimates the change in polysome vs. monosome partitioning between yeast and hyphal samples. Isoforms, isoform clusters, or genes were considered to have a significantly differential dP if there was at least a 1.5x fold change between yeast and hyphae at < 5% FDR. Based on this criterion, we defined two sets of genes, isoforms, or isoform clusters with morphology dependent dP for yeast or hyphae respectively. For each of the morphology specific gene sets, we tested for significant enrichment of the equivalent morphology specific isoform or isoform cluster set using Fisher’s exact test.

## Acknowledgements

This work was supported by the Sandler Program For Breakthrough Biomedical Research - Independent Postdoctoral Fellow Research Award (to MCK), and NIH 5R37AI066224 (to AS). AS is a BioHub San Francisco Investigator. We thank Dr. Erica Hutchins (UCSF, Department of Cell and Tissue Biology) for use of equipment for sucrose gradient preparation and fractionation. We acknowledge SeqCenter (Pittsburgh, PA), and UCSF CAT (supported by UCSF PBBR, RRP IMIA, and NIH 1S10OD028511-01 grants) for RNA sequencing. We acknowledge UCSF PCAT for use of equipment. We thank the members of the Sil and Noble labs for helpful discussions.

---

**Figure S1.**
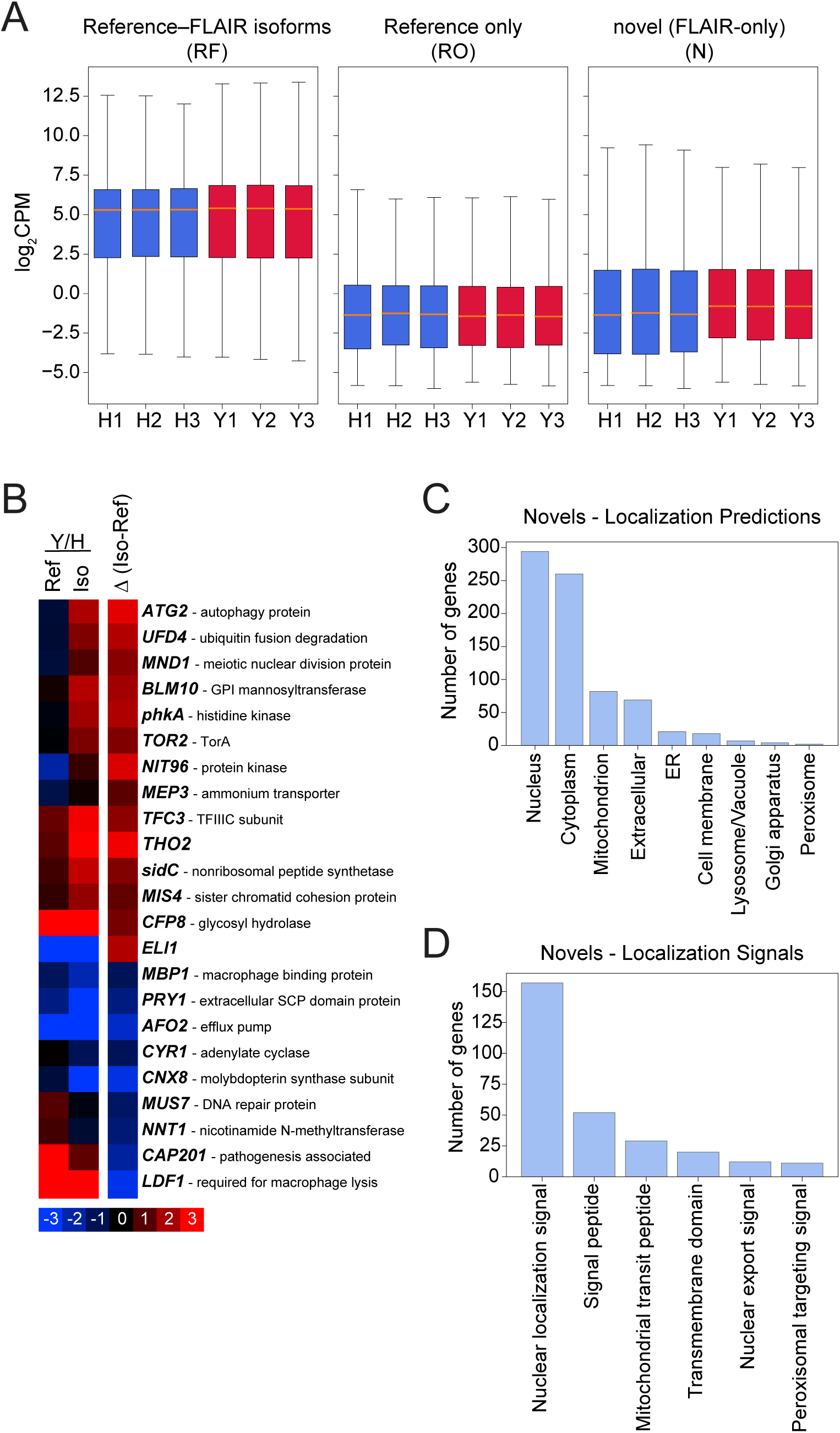
Features of isoforms and genes in each assembly category. A) Abundance (log_2_CPM) of genes that fall under RF, RO and N categories for each independent replicate. B) Comparison of Y/H differential expression using old reference gene models (Ref) and our new isoform models (Iso) for select genes with high divergence between two differential expression estimates. Δ(Iso-Ref) shows the difference of log_2_(Y/H) differential expression values. C-D) Localization and localization signal prediction using DeepLoc2.

**Figure S2.**
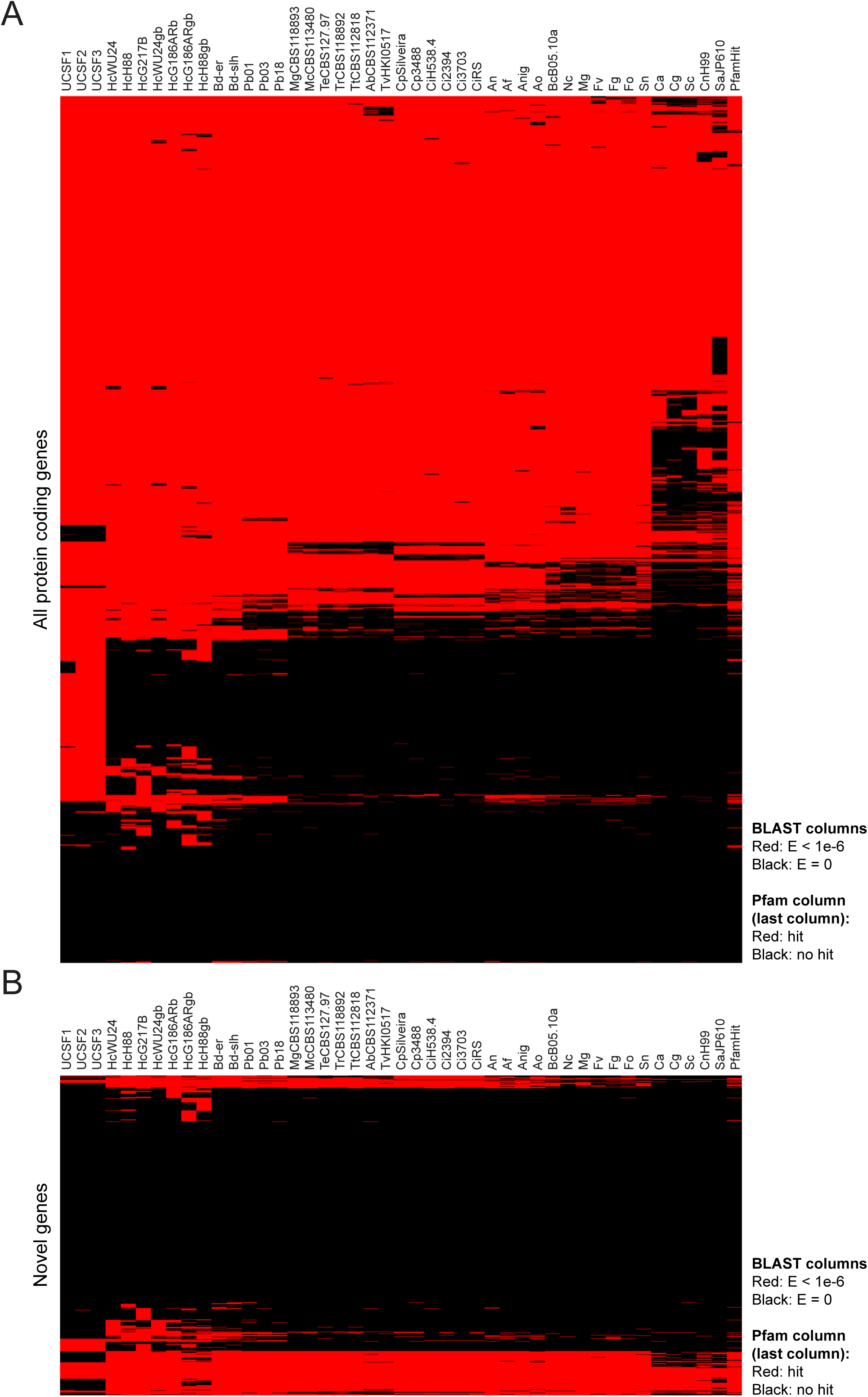
BLASTP analysis of all protein coding genes and novel genes. A) BLASTP analysis of all protein coding genes using the largest open reading frame. BLASTP analysis included other *Histoplasma* strains and species, as well as other relevant fungal organisms. Presence of any PFAM domain in the query is shown in the last column (PfamHit). B) BLASTP analysis of novel genes.

**Figure S3.**
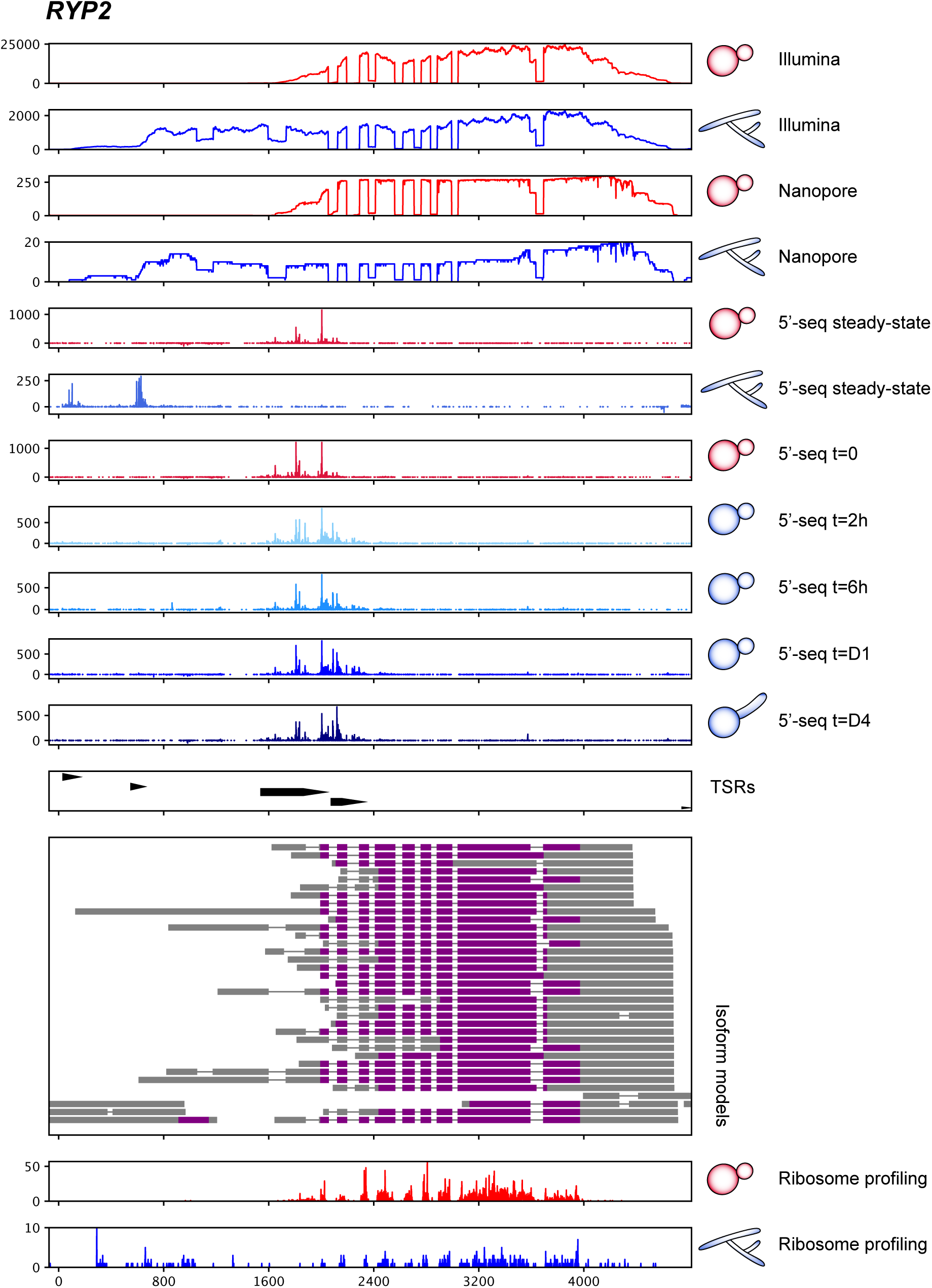
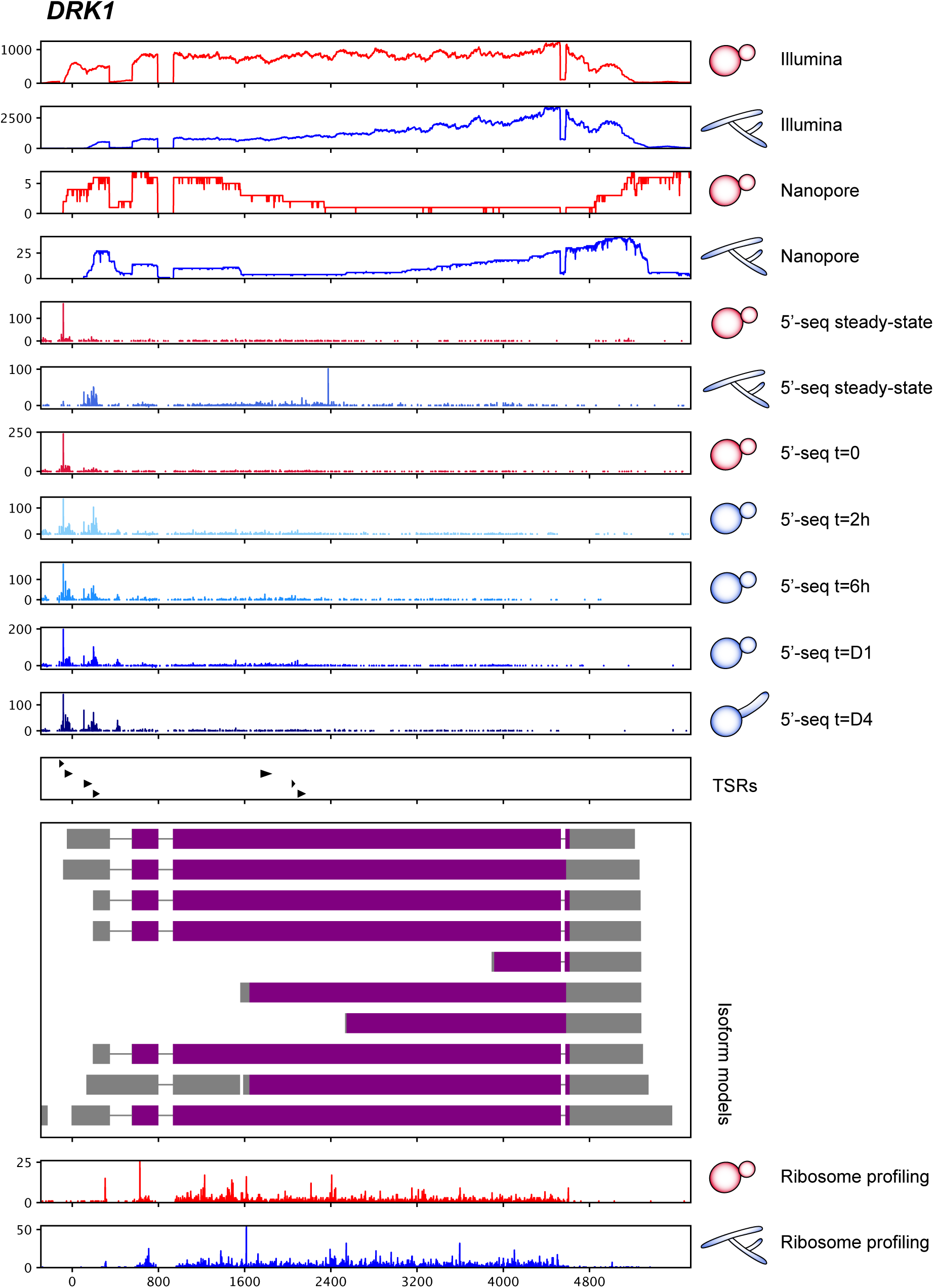
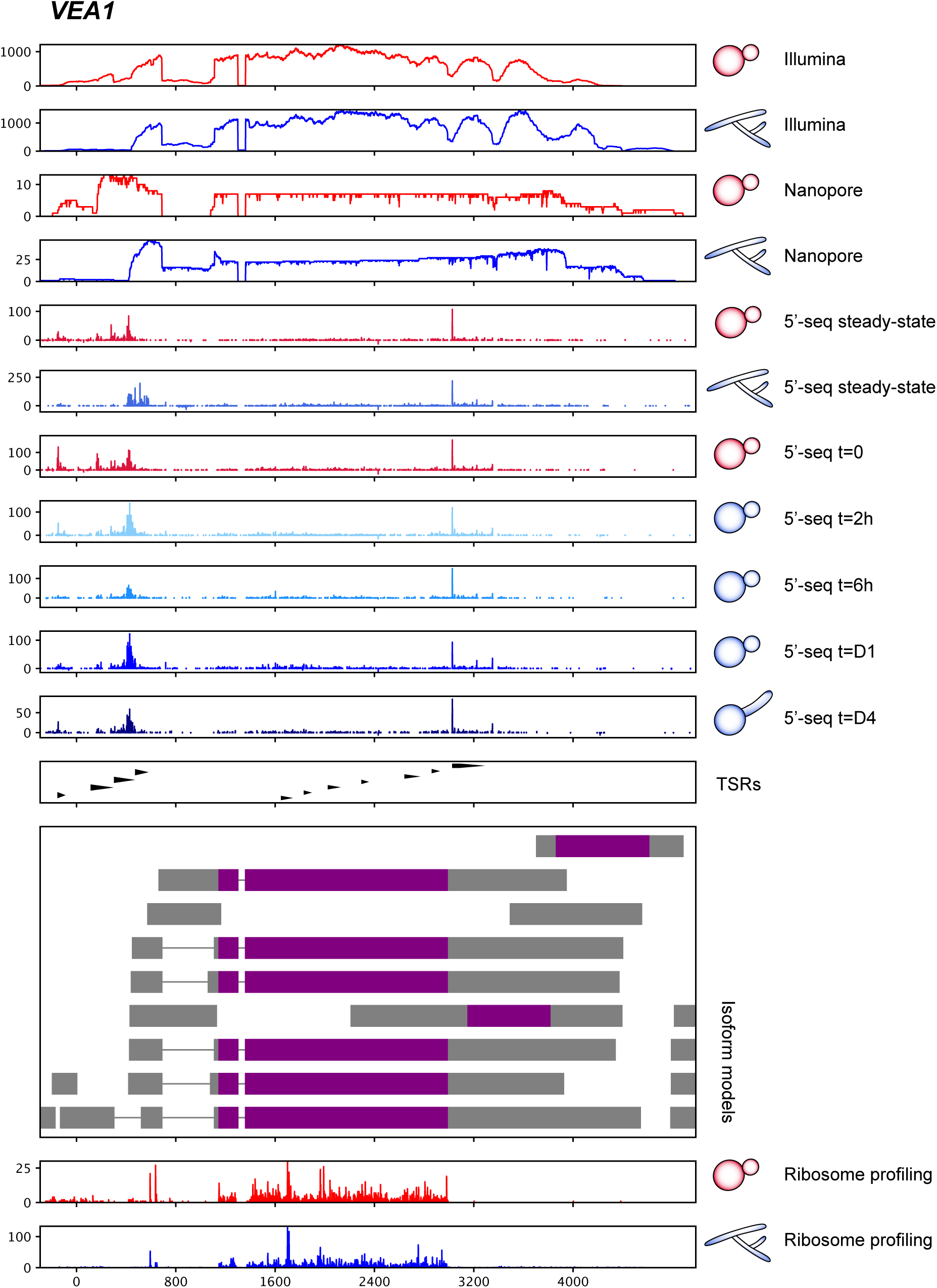
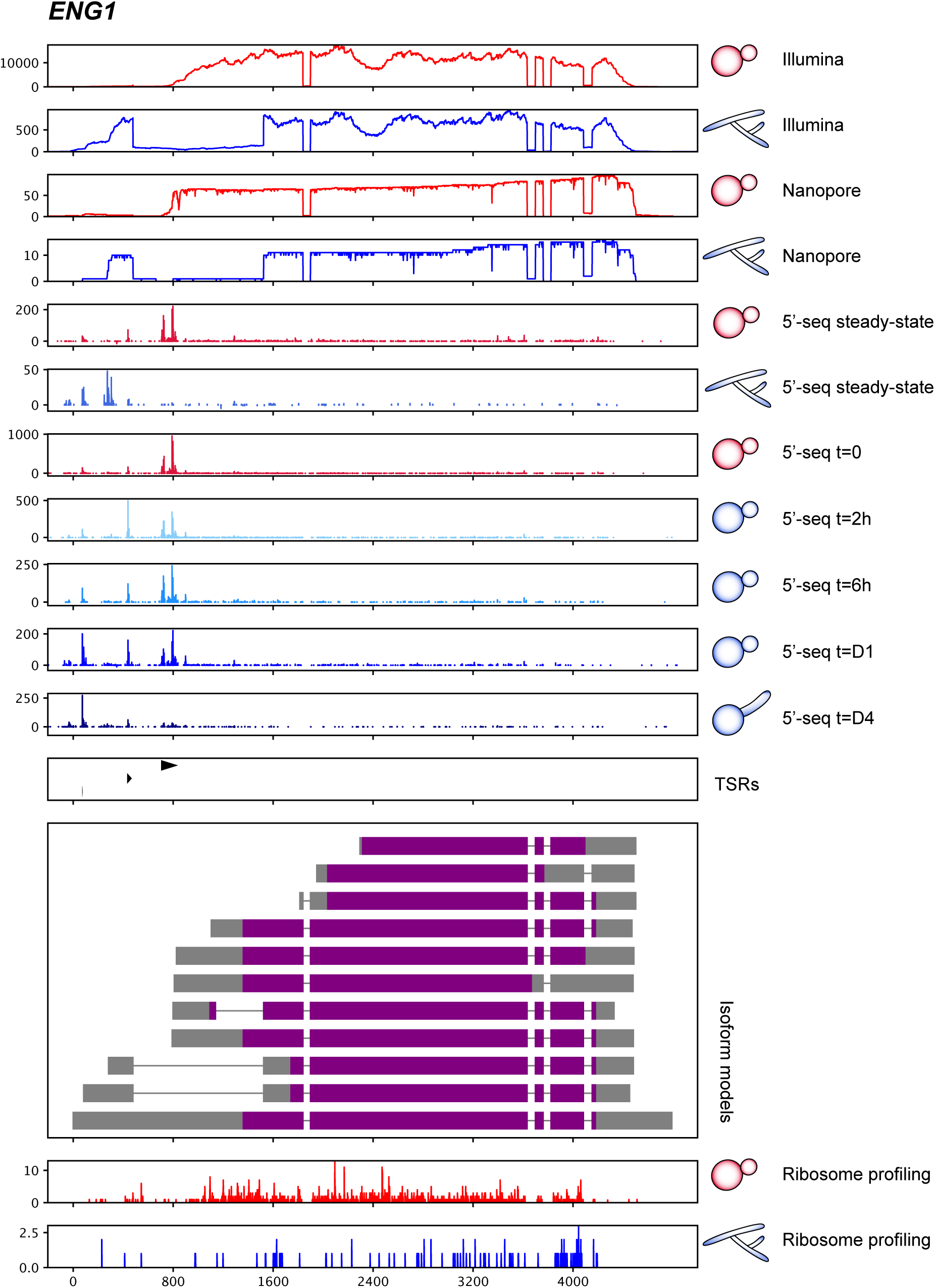
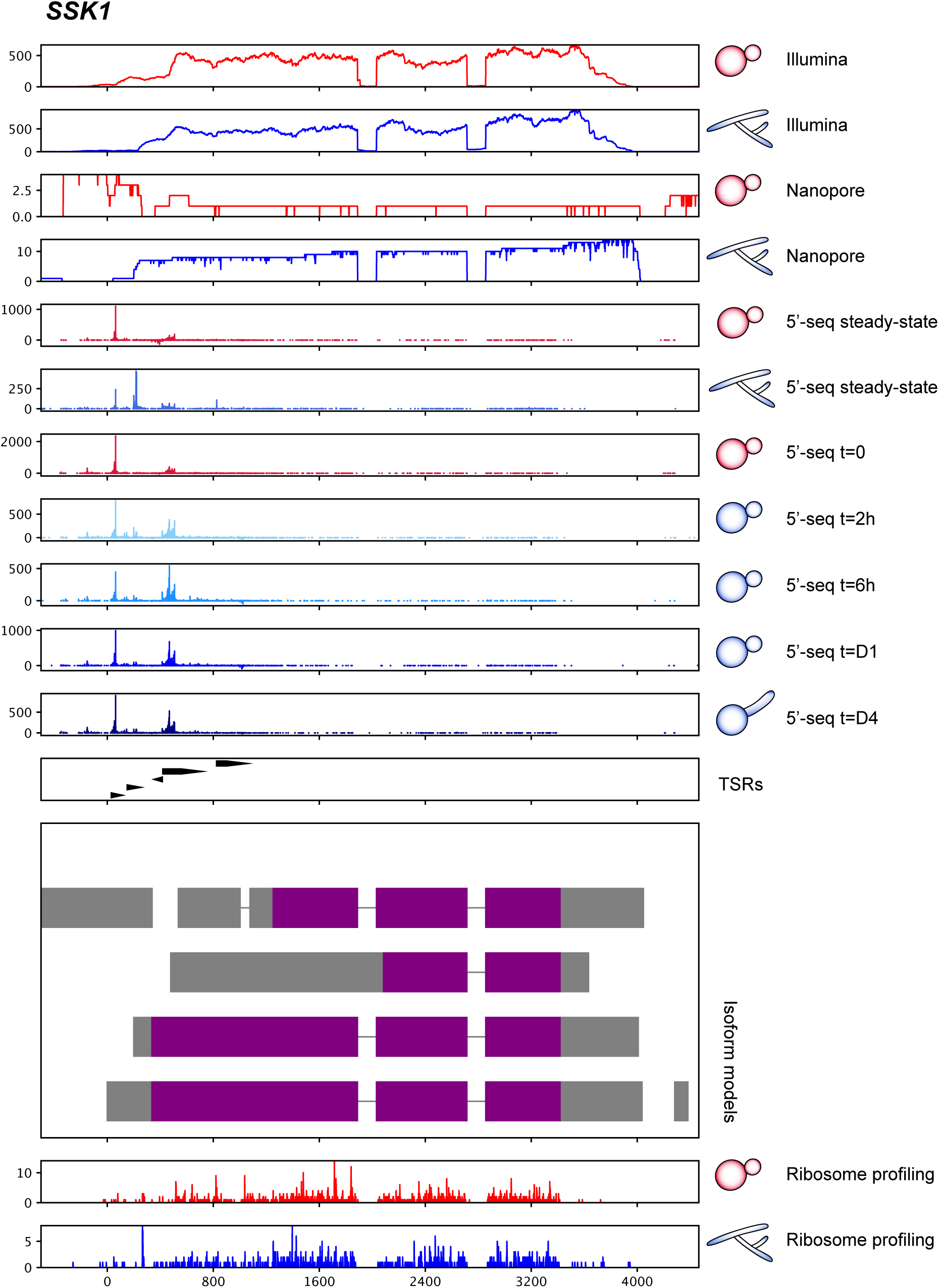
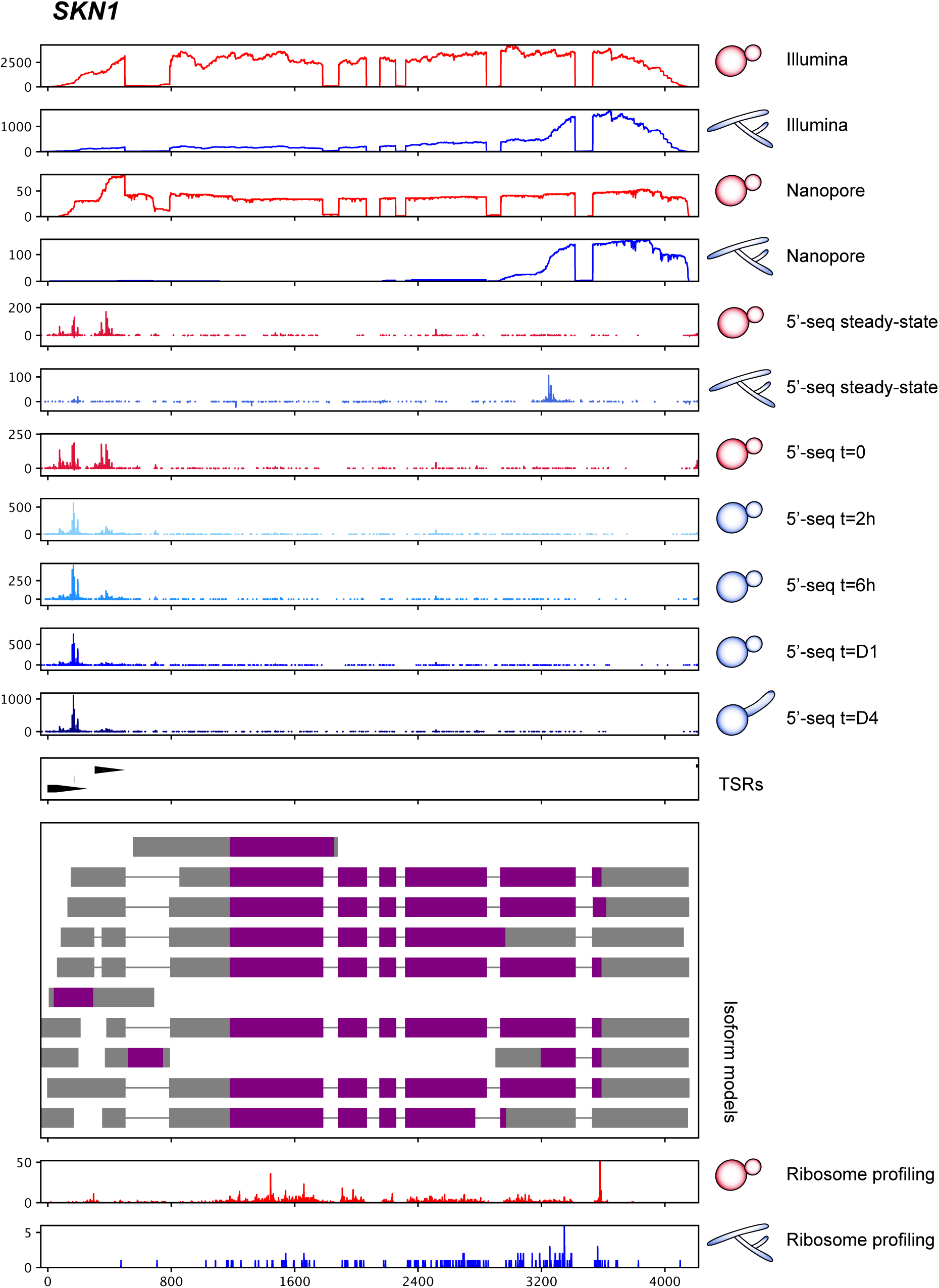
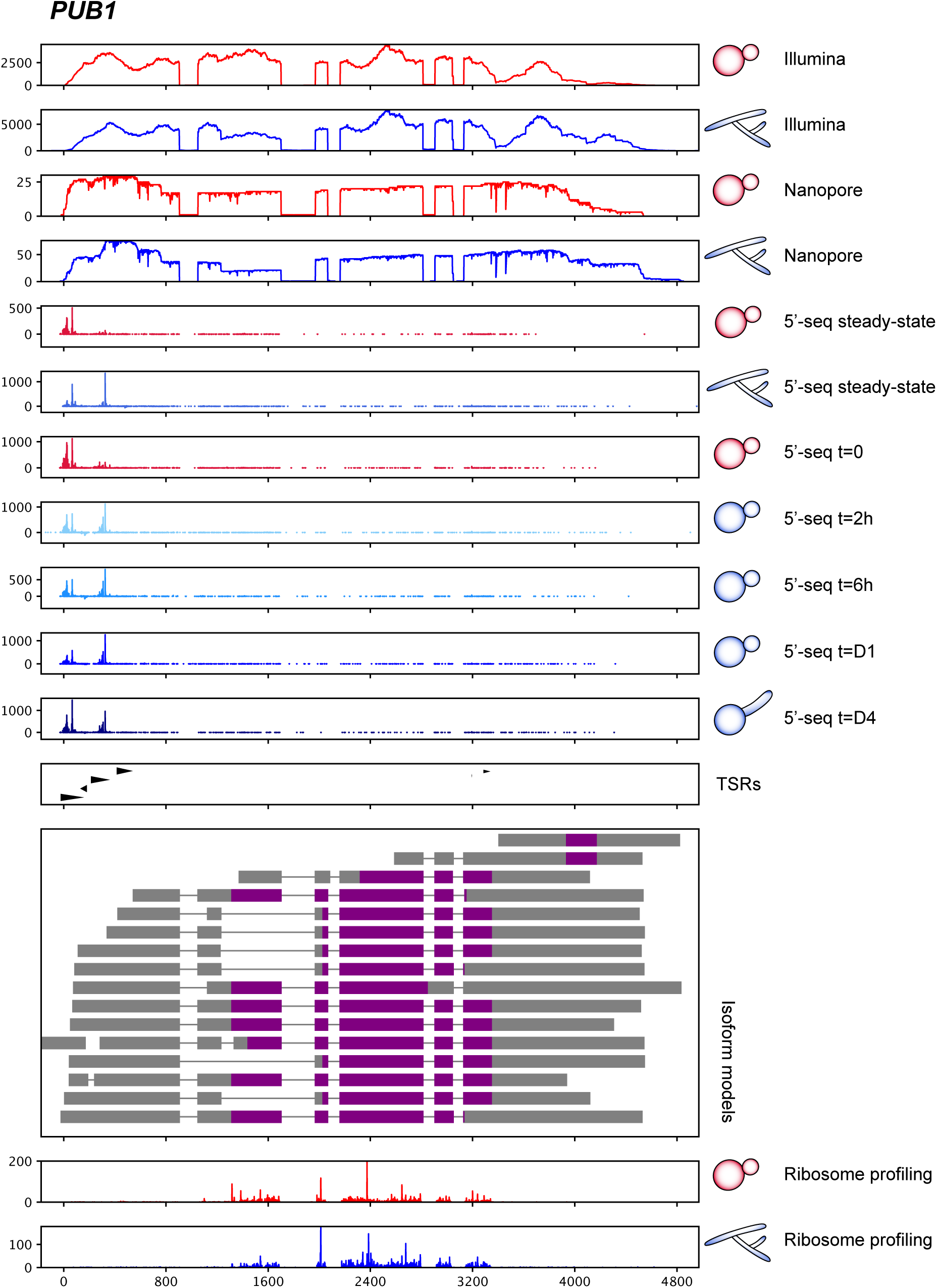
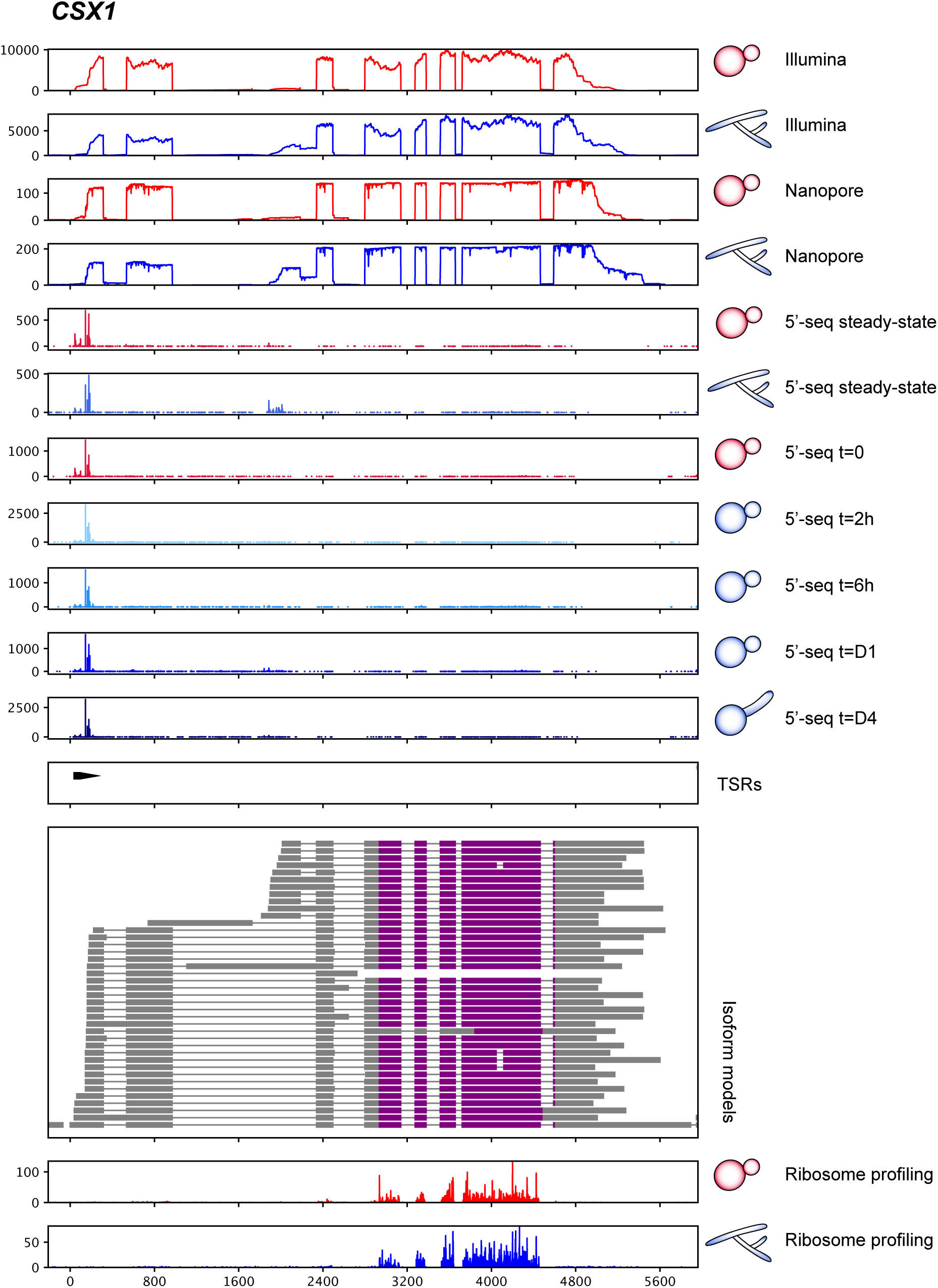
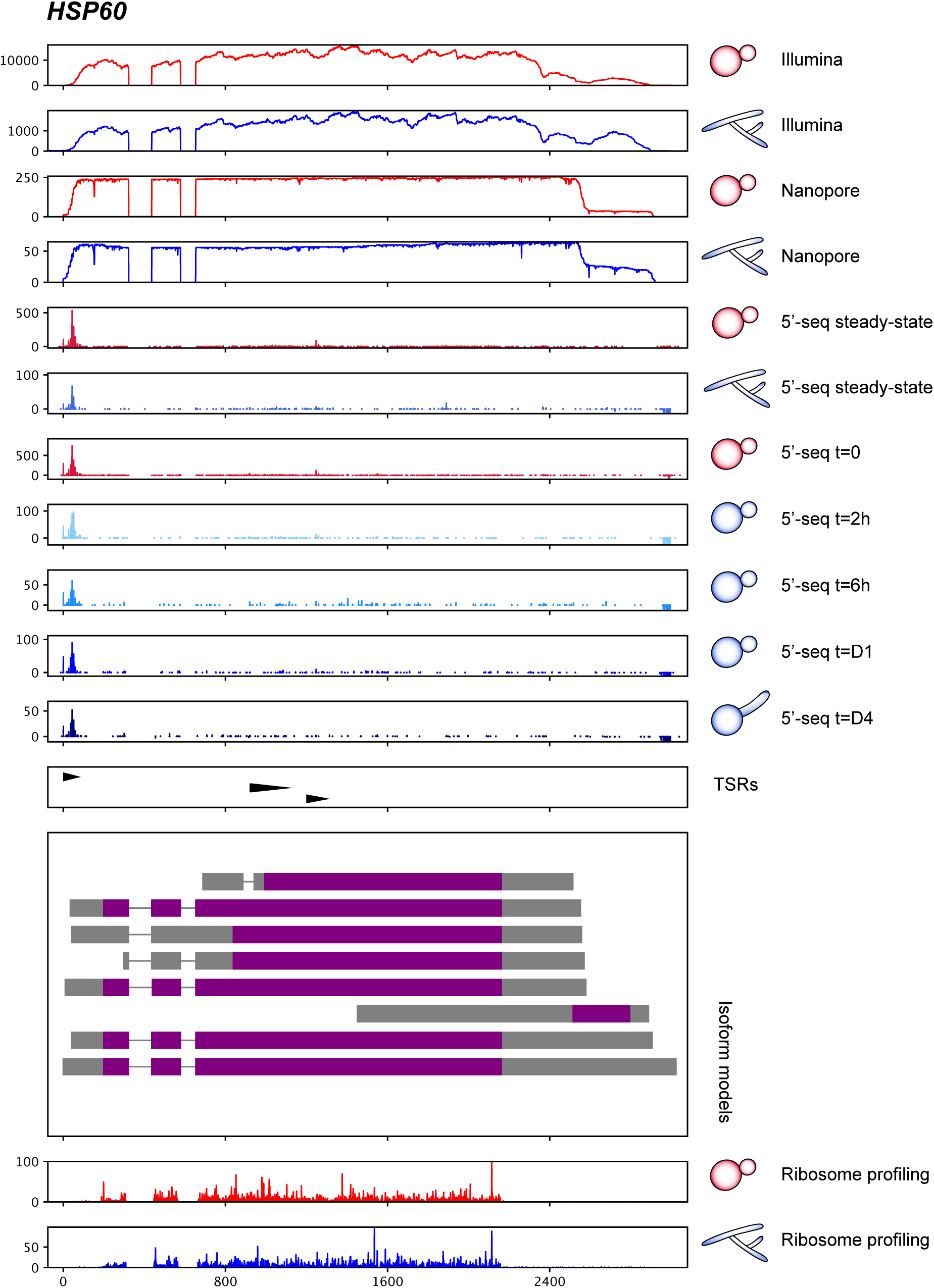
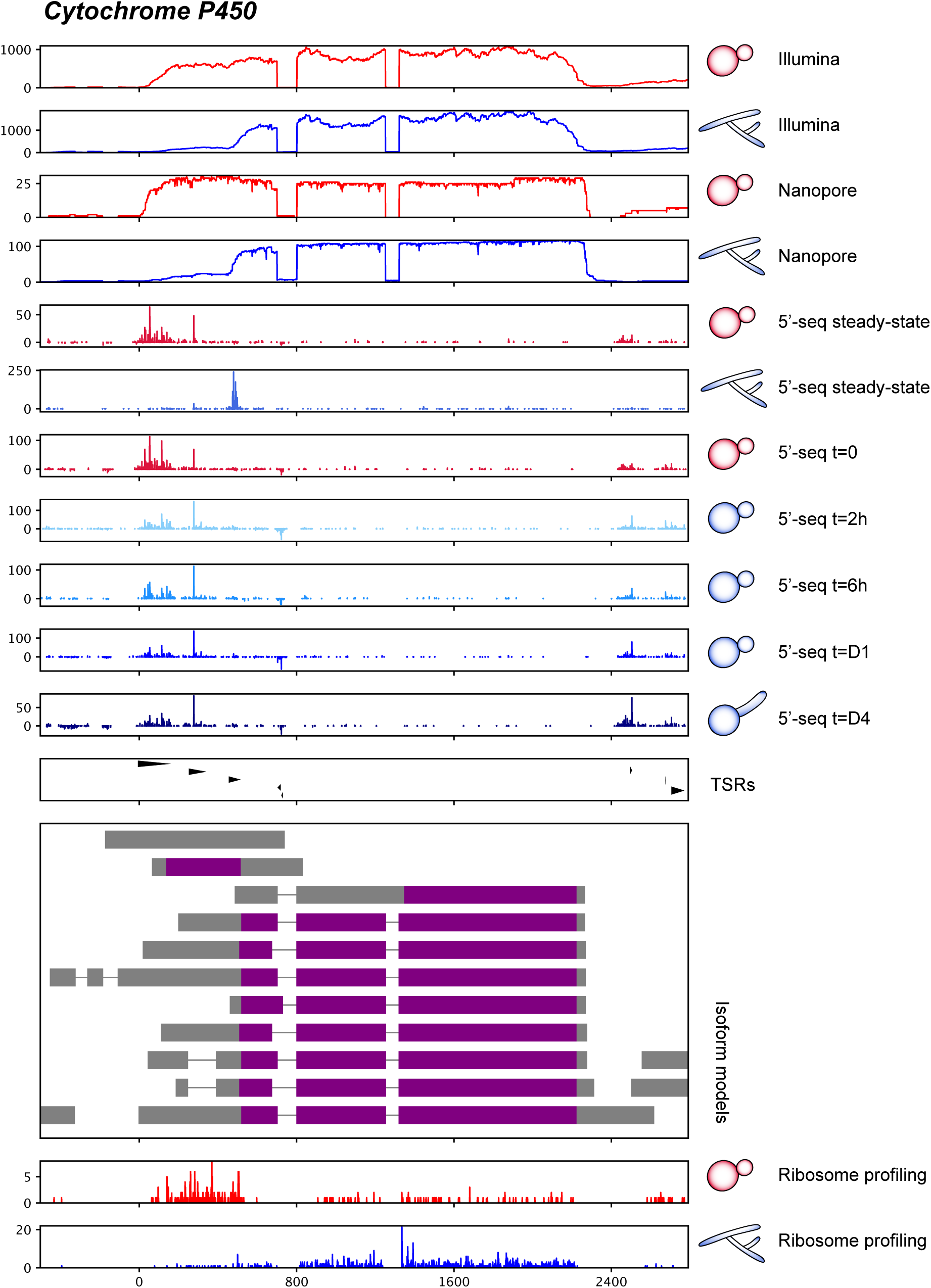
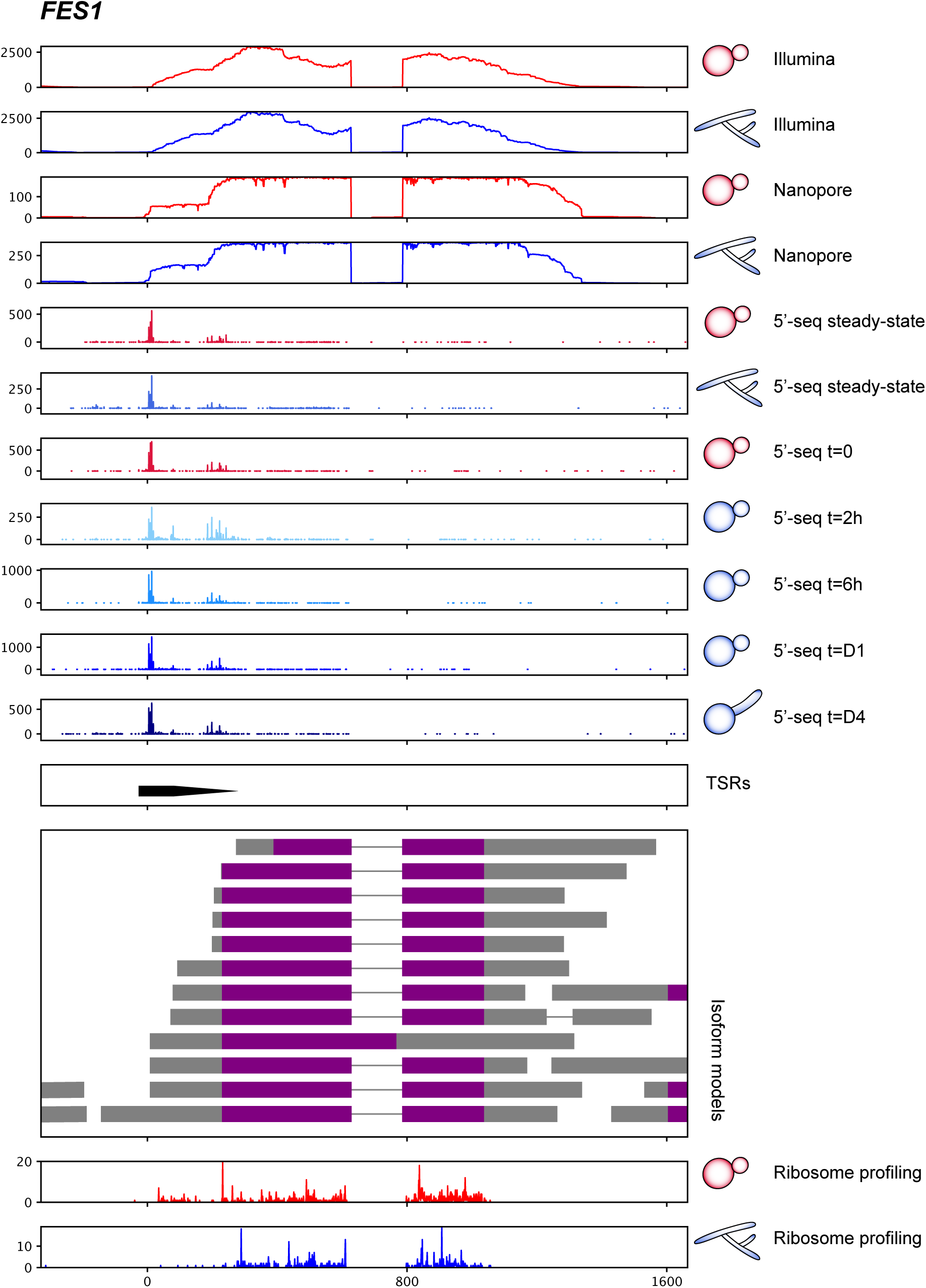
Comprehensive genome browsers. Different datasets and analyses are highlighted in an inclusive browser view for genes presented throughout the main text. Genes include *RYP2* (A), *DRK1* (B), *VEA1* (C), *ENG1* (D), *SSK1* (E), *SKN1* (F), *PUB1* (G), *CSX1* (H), *HSP60* (I), *Cytochrome P450* (J), *FES1* (K). Tracks 1-2: Illumina RNA-seq read coverage for steady-state yeast and hyphae. Tracks 3-4: 5’-adapter filtered Nanopore direct RNA-seq read coverage for steady-state yeast and hyphae. Tracks 5-6: 5’-seq data for steady-state yeast and hyphae. Tracks 7-11: 5’-seq data for 37°C to 22°C temperature shift time course. Track 12: TSR annotations. Track 13: Final FLAIR isoform models with CDS highlighted in purple. Track 14-15: Ribo-seq data as read counts per inferred ribosome P-site location for yeast and hyphae.

**Figure S4.**
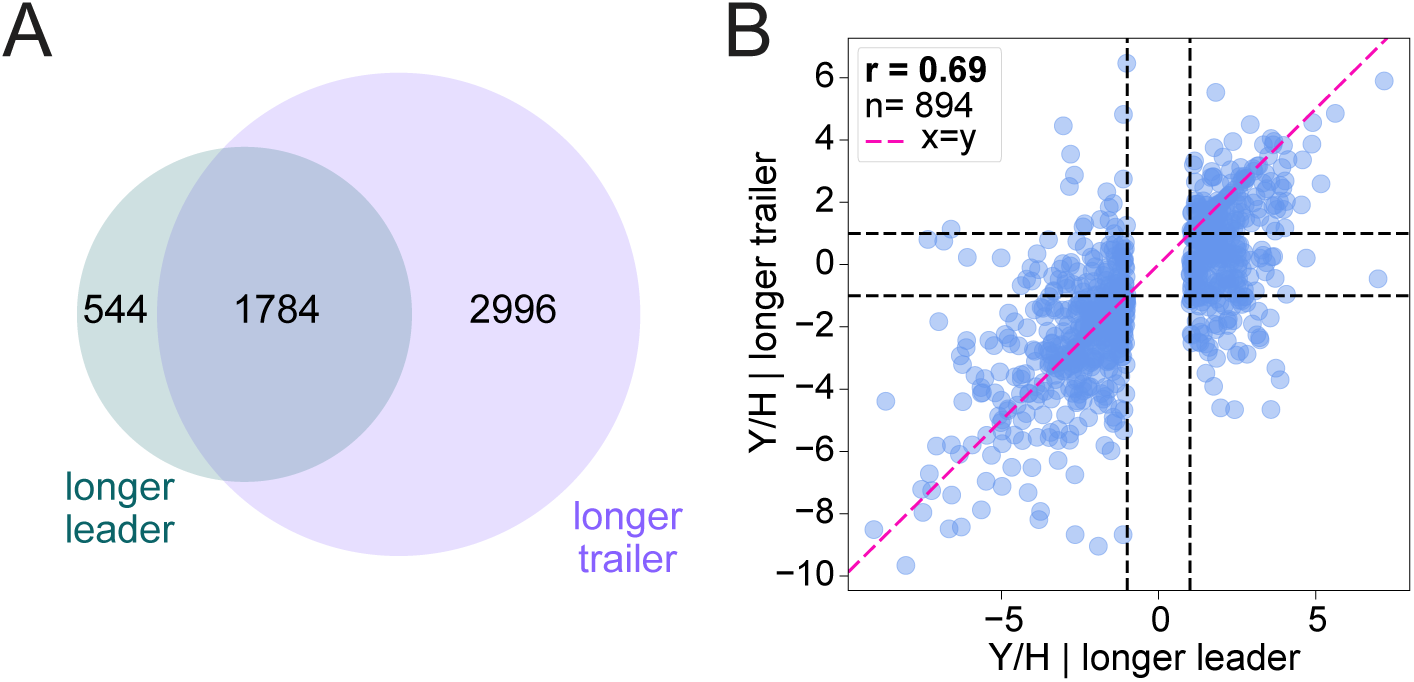
Some genes encode both long 5’ leader and long 3’ trailer isoforms. A) Venn diagram of genes with long 5’ leader and long 3’ trailer isoforms. B) Scatter plot shows the correlation between the log_2_(Y/H) expression of long 5’ leader isoforms vs. long 3’ trailer isoforms of genes whose long 5’ leader isoforms are significantly differentially expressed (2-fold cutoff and 5% FDR). The pink line represents x = y. Pearson correlation coefficient is highlighted. “n” values indicate the number of genes. Dashed black lines represent a 2-fold change.

**Figure S5.**
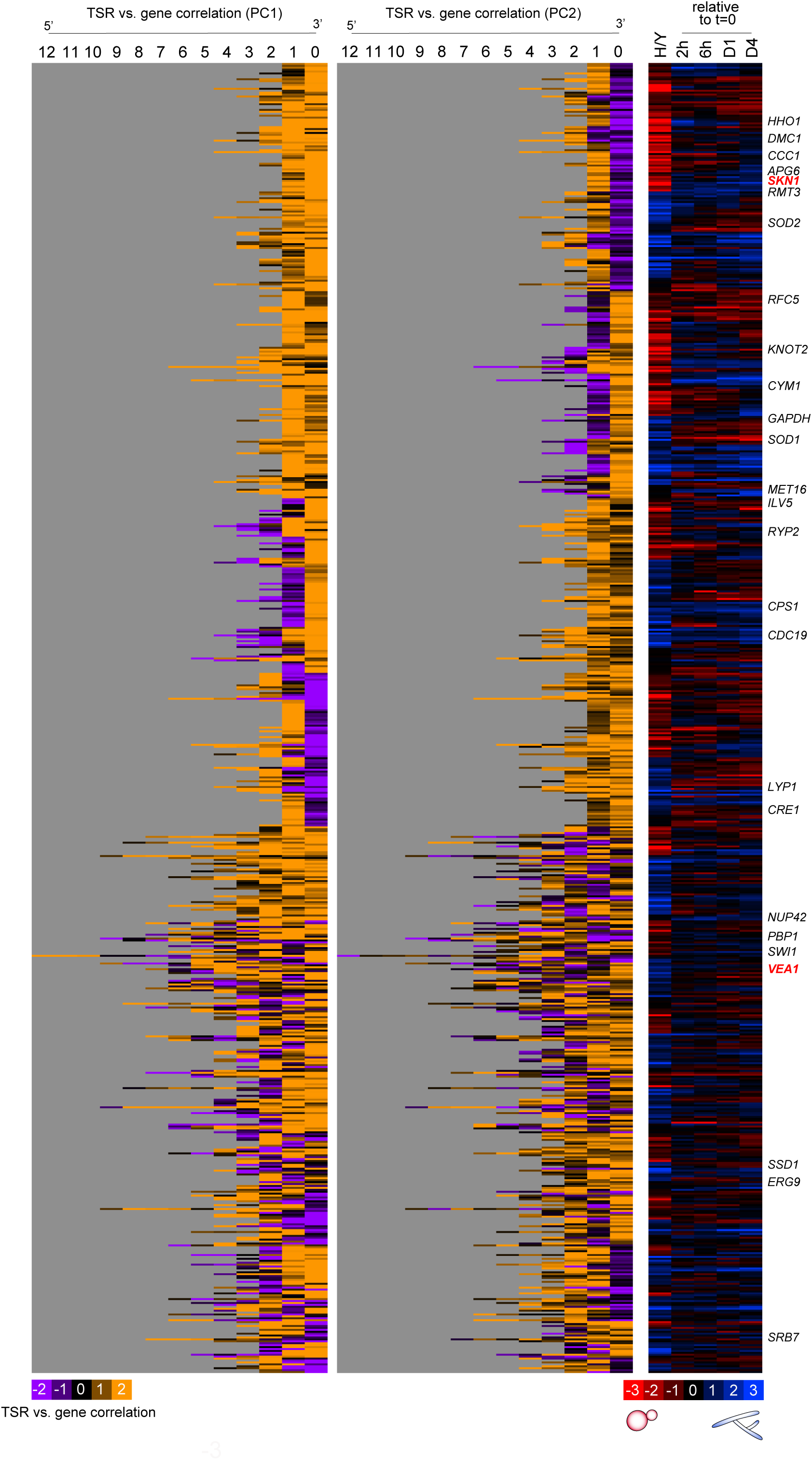
Temperature-responsive TSR expression changes (class 2). Heatmaps show genes where two principal components (PC1 and PC2) explain the variance across TSRs of a given gene (class 2 genes). *Right panel (red-blue):* Differential expression of all summed TSRs of a given gene in steady state hyphae relative to yeast (H/Y), or relative to mid-log yeast at 37°C (t = 0) across time points. *Left panels (orange-purple):* PC1 and PC2 weights on each TSR showing TSR to gene correlation. PCs are oriented so that positive weight (orange) indicates correlation with the gene level expression pattern and negative weight (purple) indicates anticorrelation. TSR 0 denotes the most 3’ TSR and TSR 9 denotes the most 5’ TSR relative to the start codon. Rows are clustered by 5’ to 3’ ordering of PC1 weights on TSRs, then by gene-level expression pattern. Gray indicates no data.

**Figure S6.**
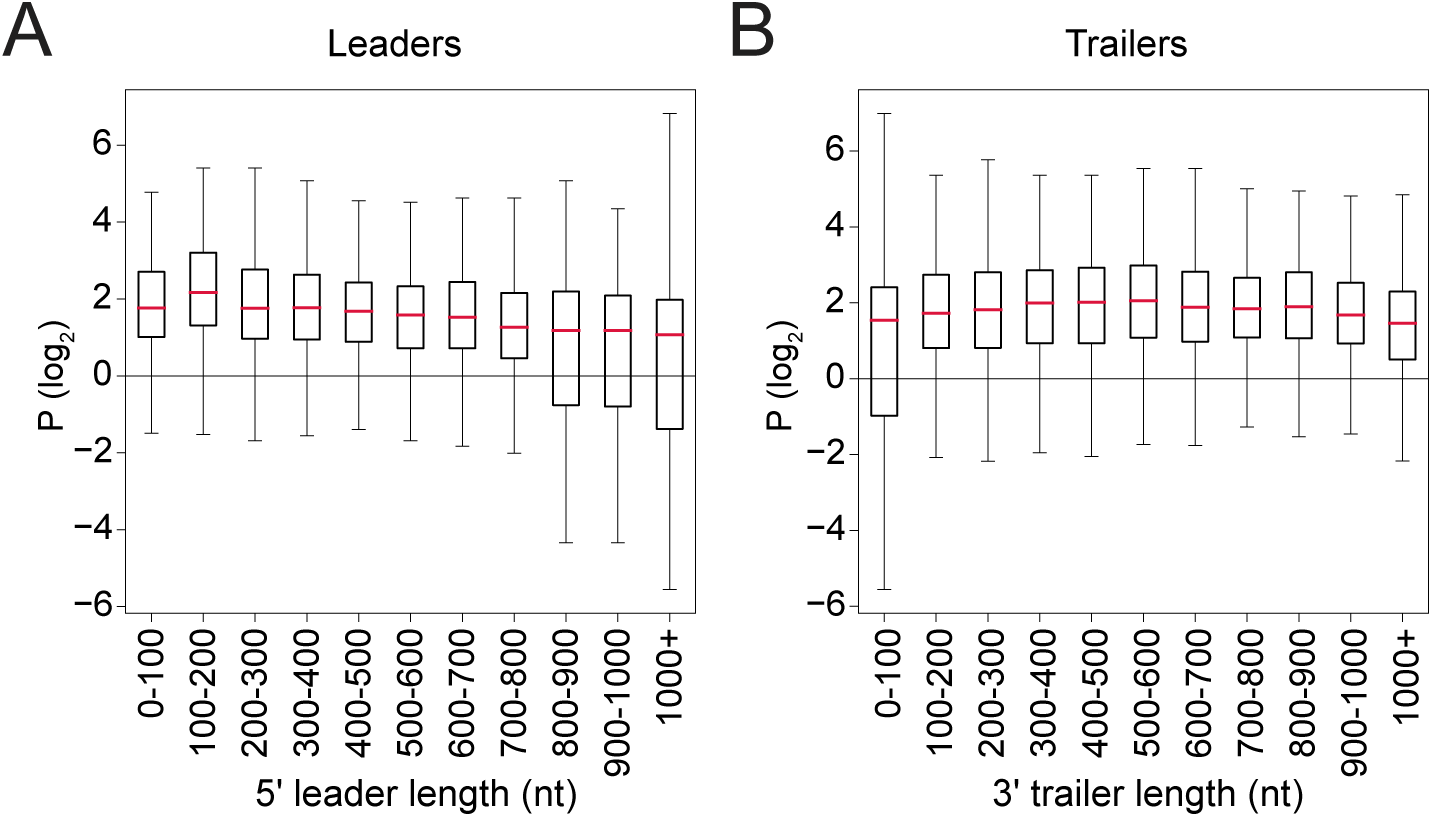
Polysome association of all isoforms grouped by 5’ leader and 3’ trailer length. A) Box plots showing polysome association (P) of all isoforms grouped by 5’ leader length bin. Only isoforms of genes encoding both long and short 5’ leader isoforms are included. Red bars indicate the median, and boxes indicate interquartile range (IQR). B) Box plots showing P for all isoforms grouped by 3’ trailer length bin.

**Figure S7.**
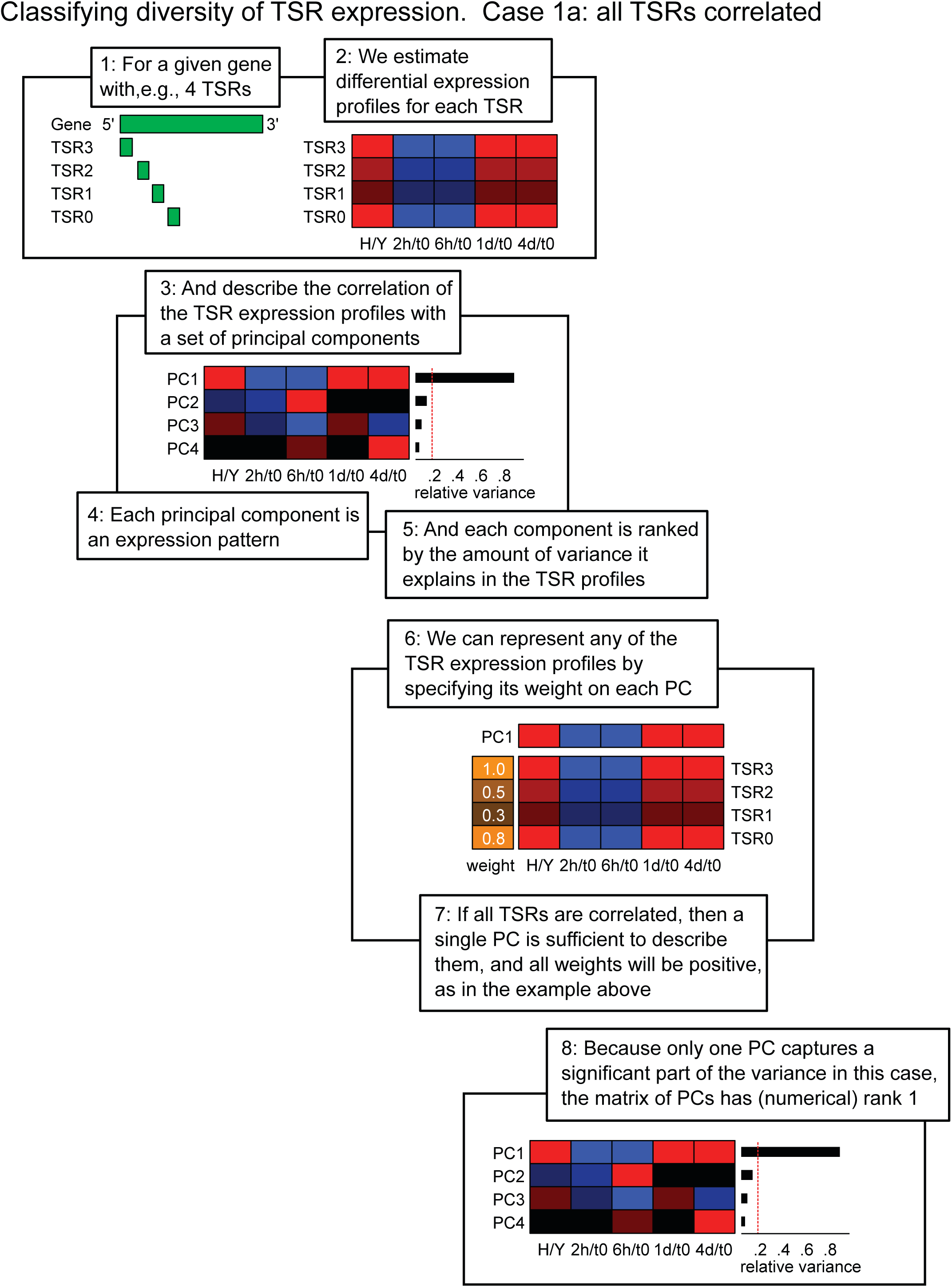
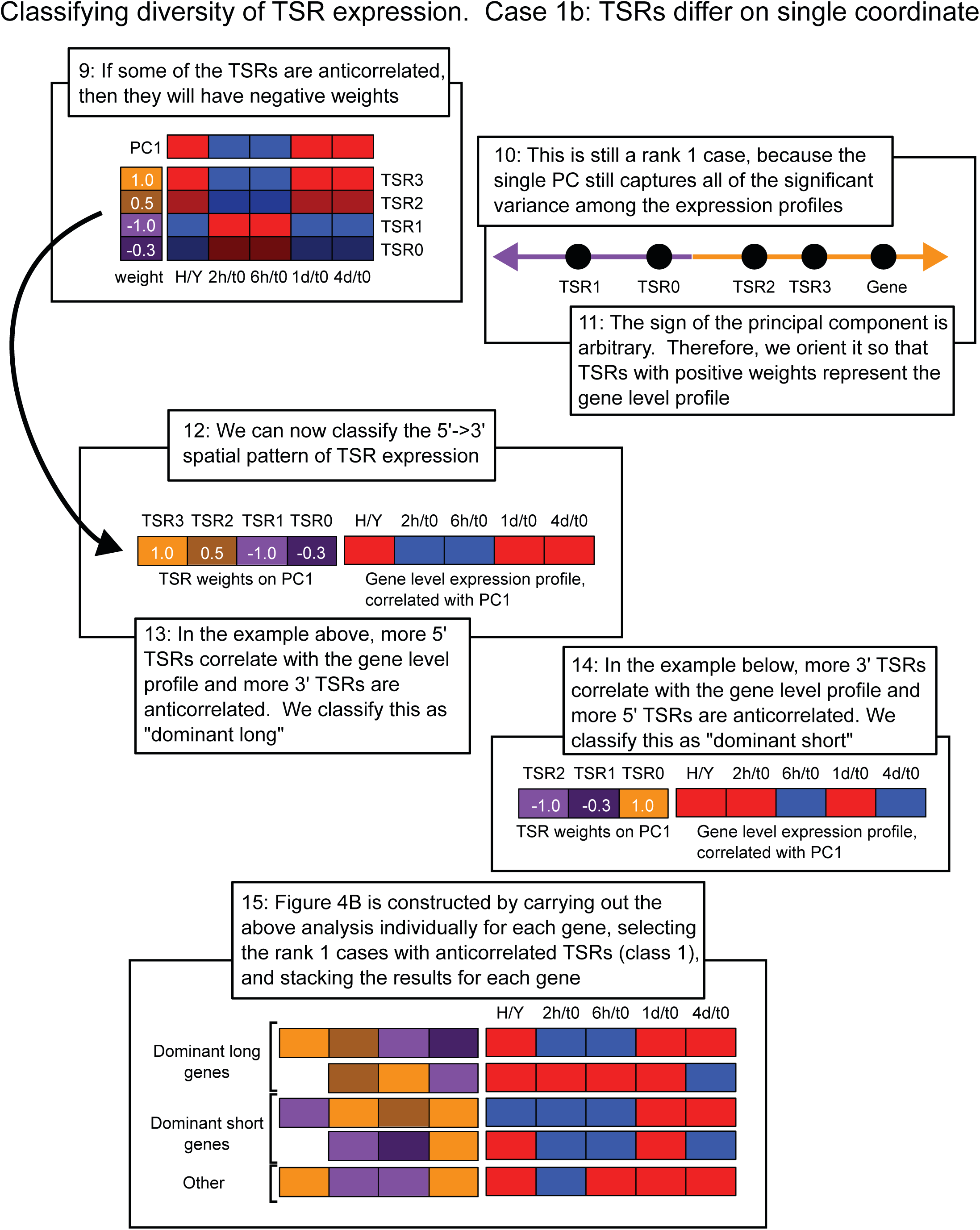
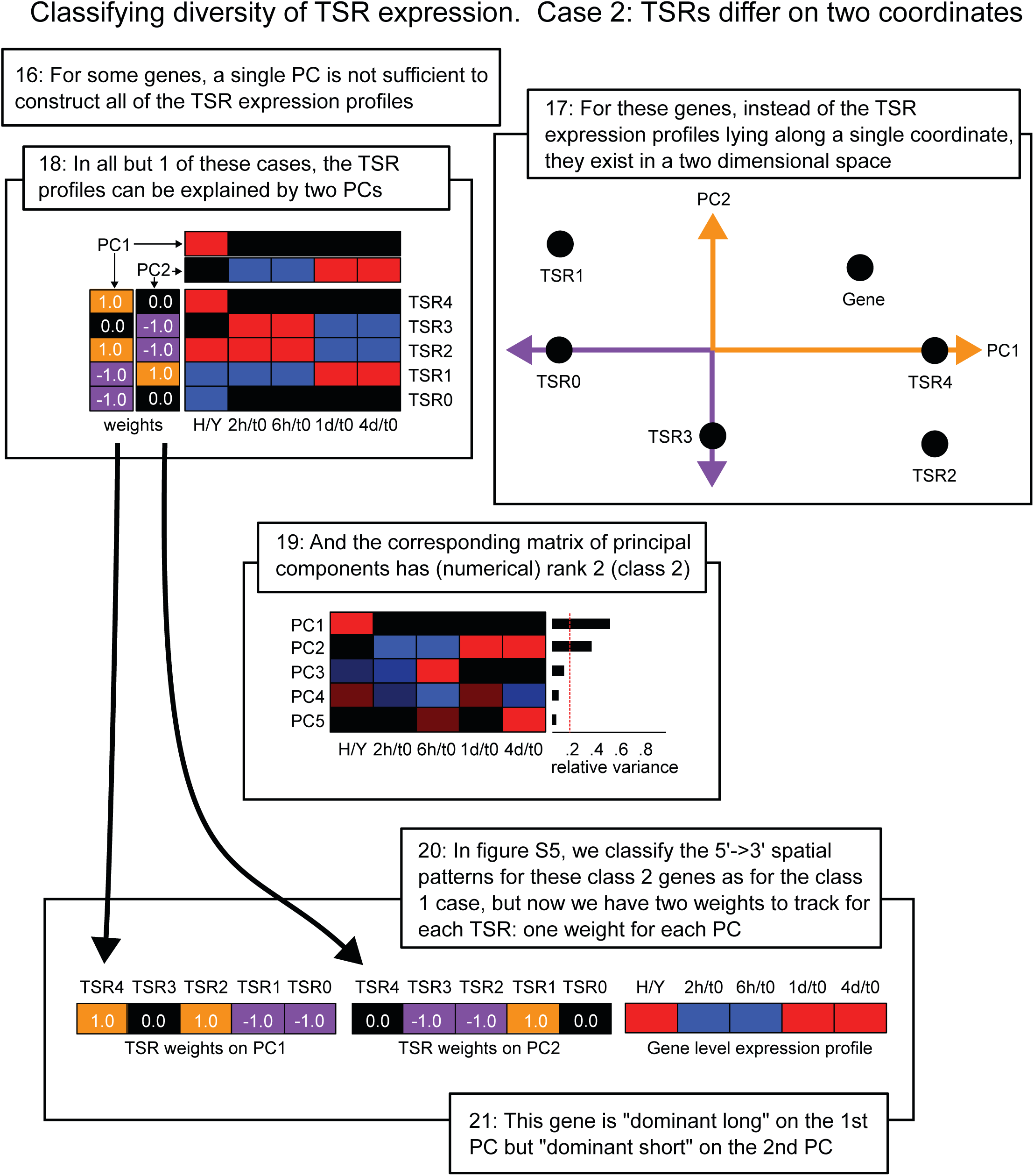
Classification of TSR expression profiles post-temperature shift from 37°C to 22°C using PCA analysis. Summary of PCA analysis steps to determine TSR expression complexity.

